# Structure of CPC-PP2A(B56) Kinase-Phosphatase Signaling Hub in Chromosome Segregation Fidelity

**DOI:** 10.64898/2026.11.18.767505

**Authors:** Paula Sotelo-Parrilla, Vimal Das, Ryu-Suke Nozawa, Maria A. Abad, Pratiti Nanda, Zhaoyue Yan, Zephyr Ford, Juan Zou, Christos Spanos, Lucia Fernández Arnáez, Juri Rappsilber, Toru Hirota, A. Arockia Jeyaprakash

**Affiliations:** Gene Center Munich, Ludwig-Maximilians-Universität Munchen, Munich, Germany; Centre for Cell Biology, University of Edinburgh, Edinburgh, UK; Division of Experimental Pathology, Cancer Institute of the Japanese Foundation for Cancer Research, Tokyo, Japan; Institute of Biotechnology, Technische Universität Berlin, 13355 Berlin, Germany

## Abstract

Accurate chromosome segregation relies on kinase-phosphatase crosstalk that controls protein interactions underpinning sister chromatid cohesion, error correction and timely anaphase onset. The PP2A(B56) phosphatase and the Aurora B kinase-containing Chromosomal Passenger Complex (CPC), two central mitotic regulators, are independently recruited to centromeres by Shugoshin1 (Sgo1). Here, using biochemical reconstitutions and cryo-electron microscopy, we show that PP2A(B56) and CPC form an integrated signaling platform independent of Sgo1. The CPC binds PP2A(B56) directly through two Borealin elements: a previously uncharacterized LxxIxE-like motif and the dimerization domain. Sgo1 selectively remodels the dimerization-domain interface, while the LxxIxE-like motif sustains PP2A(B56)-CPC association. Disrupting this interaction increases chromosome segregation errors, compromising segregation fidelity. Our findings reveal a direct kinase-phosphatase association as an additional layer of mitotic regulation that safeguards segregation fidelity.

## Introduction

Accurate chromosome segregation during mitosis is fundamental to genomic stability. Errors in this process can lead to aneuploidy, a hallmark of cancer and developmental disorders. Faithful chromosome segregation requires biorientation, mediated by the physical attachment of sister chromatid kinetochores to microtubules emanating from opposite spindle poles (*1–7*). The kinetochore is a multi-subunit protein structure that assembles at the centromere, a chromosomal region characterized by the enrichment of CENP-A containing nucleosomes (*8–11*).

In addition to serving as the mechanical interface between the chromosomes and spindle microtubules, centromeres function as signaling hubs where kinase and phosphatase activities are precisely coordinated in space and time to tune protein-protein interactions. This ensures the destabilization of incorrect kinetochore-microtubule attachments and the selective stabilization of the correct ones (a process known as error correction), and the maintenance of sister chromatid cohesion until the spindle assembly checkpoint (a quality control mechanism that monitors attachments and delays anaphase onset until biorientation is achieved) is silenced (*12–14*). In particular, the Chromosomal Passenger Complex (CPC; composed of Borealin, Survivin, INCENP and Aurora B kinase) and protein phosphatase 2A containing the B56 regulatory subunit (PP2A(B56)) form a key kinase-phosphatase pair whose dynamic crosstalk coordinates all these processes (*14*).

PP2A is a trimeric Ser/Thr phosphatase consisting of a catalytic (C) and scaffolding (A) subunit core that associates with one of four families of regulatory (B) subunits responsible for substrate specificity. The B56 family, containing isoforms α, β, γ, δ and ε, is characterized by the presence of a highly conserved pocket that recognizes proteins containing LxxIxE-like motifs (*15–18*). B56 isoforms localize to distinct chromosomal sites via specific recruiters (*19*), with PP2A(B56α) and PP2A(B56ε) predominantly targeted to inner centromeres through their interaction with Shugoshins 1 and 2 (Sgo1 and Sgo2); and PP2A(B56γ) and PP2A(B56δ) enriched at kinetochores via interaction with BubR1 and Sgo1 (*19*). CPC exerts distinct roles during mitosis as it dynamically transitions between chromosome arms, centromeres, the central spindle, and the midbody. Its centromeric enrichment in prometaphase/metaphase is driven by multiple recruitment pathways, including Haspin- and Bub1-mediated histone phosphorylations that recruit Survivin and Sgo1, respectively (20–33); as well as histone phosphorylation-independent mechanisms involving Borealin and INCENP-mediated interaction with histones and nucleosomal DNA (30, 34–36).

Emerging evidence suggests that the coordinated integration of opposing kinase and phosphatase activities, mediated by shared recruiting factors, is critical for accurate chromosome segregation (*37–39*). Simultaneous recruitment of kinase-phosphatase pairs directly influences the local phosphorylation/dephosphorylation turnover of kinetochore and centromeric substrates (*14, 37–39*). For both CPC and PP2A(B56), Sgo1 serves as a molecular scaffold required for their centromeric recruitment. The Sgo1 N-terminal region establishes direct interactions with Survivin and Borealin (*27, 33, 40, 41*), and the Sgo1 N-terminal dimerization helices (Sgo1_DH)_ engage PP2A B56 and C subunits (*42*). However, whether Sgo1 can interact with CPC and PP2A(B56) simultaneously to assemble an integrated kinase-phosphatase signaling platform, and, if so, the structural basis and functional requirement of such assembly for ensuring chromosome segregation fidelity, remain critical open questions.

Here, using an integrated biochemical, structural and cell biological approach, we demonstrate that Sgo1 can simultaneously engage both PP2A(B56) and CPC, and provide the structural basis for this engagement. Even more strikingly, our work reveals a direct Sgo1-independent interaction between CPC and PP2A(B56) mediated by a previously uncharacterized LxxIxE-like motif in Borealin that binds the LxxIxE-binding pocket of PP2A(B56). Disrupting this interaction increases chromosome congression defects and chromosome segregation errors, and delays anaphase onset, compromising segregation fidelity. Together, our findings uncover a novel mechanism for coordinating CPC kinase and PP2A(B56) phosphatase activities and reveal the direct interaction between counteracting CPC and PP2A(B56) as an additional layer of regulation required to ensure faithful chromosome segregation.

## Results

### Sgo1 can interact simultaneously with PP2A(B56) and CPC, forming a robust complex

Given the close proximity of PP2A(B56) and CPC binding sites on Sgo1, we investigated whether PP2A(B56), Sgo1 and CPC could assemble into a hetero-oligomeric ternary complex *in vitro*. Size-exclusion chromatography (SEC) analysis showed complex formation between the PP2A(B56) holocomplex (composed of full-length A, B56 and C subunits), an N-terminal fragment of Sgo1 (Sgo1_1-155_) and the CPC localization module (CPC_I1-58SB_, comprising the first 58 amino acids of INCENP and full-length Survivin and Borealin). The resulting complex eluted at 10.2 ml in the size exclusion chromatography analysis, compared with 12.3 ml for PP2A(B56) and 13.3 ml for Sgo1_1-155_ alone, consistent with the formation of a higher molecular weight complex (Fig. 1, A and B).

**Figure 1:**
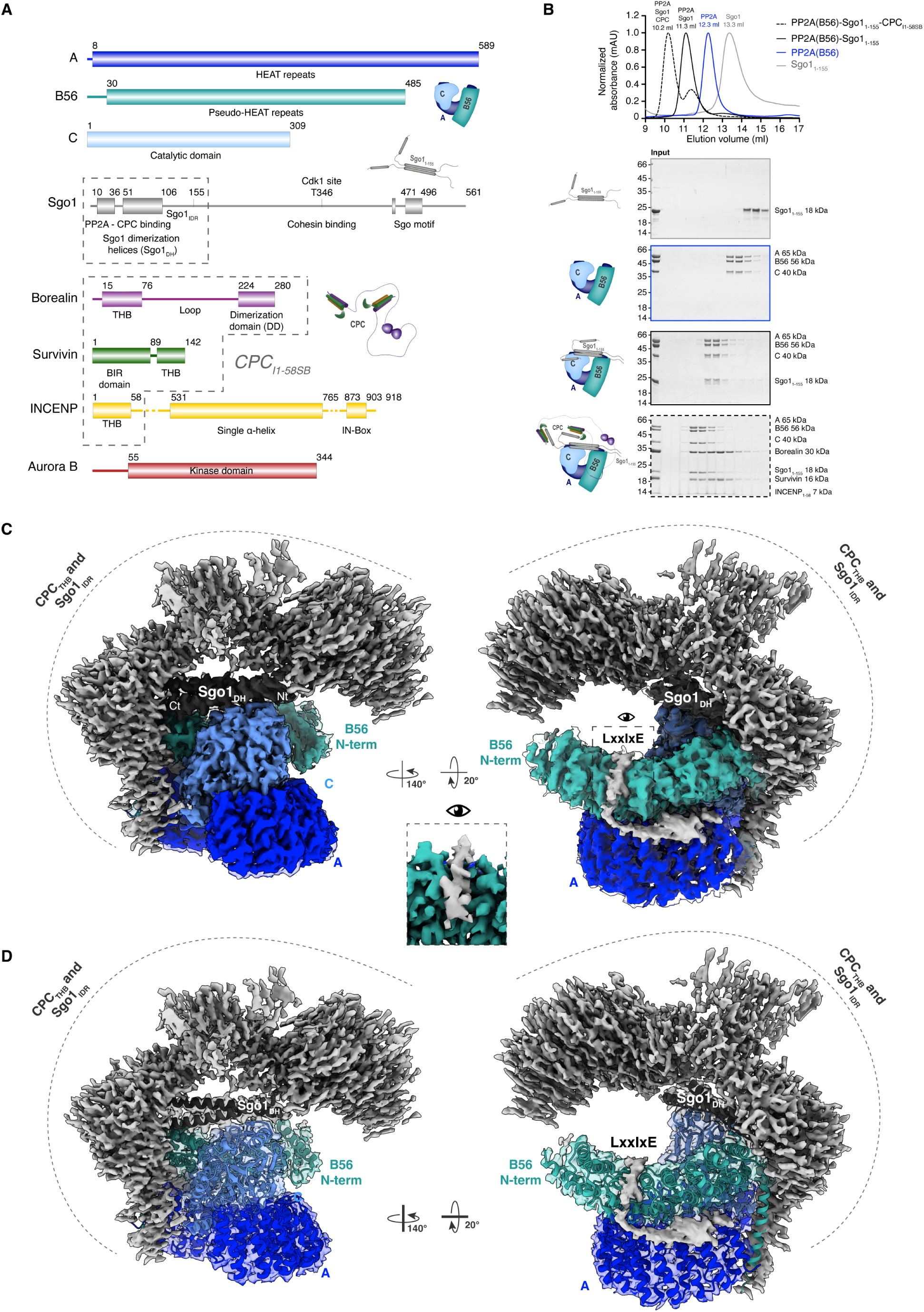
Overall assembly and architecture of the PP2A(B56)-Sgo1-CPC assembly. **(A)** Domain architecture and corresponding cartoons for PP2A(B56), Sgo1 and CPC. In all figures, A subunit is shown in blue, B56 subunit in teal, C subunit in light blue, Sgo1 in dark grey, INCENP in yellow, Survivin in green and Borealin in purple. Sgo1_1-155_ and CPC_I1-58SB_ are marked with a dashed box. The domains forming the triple helical bundle of CPC are labelled as THB (triple helical bundle). **(B)** Size Exclusion profiles, elution volume and corresponding SDS-PAGEs of Sgo11-155 (grey), PP2A(B56) (blue), PP2A(B56)-Sgo1_1-155_ (black) and PP2A(B56)-Sgo1_1-155_-CPC_I1-58SB_. Representative cartoons are shown on the left. **(C)** Overlay of the ‘consensus’, ‘high-resolution’ and ‘Borealin loop’ maps (Fig. S1). Conformationally heterogeneous densities corresponding to Sgo1 intrinsically disordered region (Sgo1_IDR_) and Sgo1-tethered CPC triple-helical bundle (CPC_THB_), as well as the additional density on the A-B56 interface, have been shown in light grey. A close-up top view of the unaccounted density observed at the PP2A(B56) LxxIxE-binding pocket is shown between the two maps. **(D)** Overlay of the cryo-EM density maps with the high-resolution map displayed with transparency and the PP2A(B56)-Sgo1 dimerization helices (Sgo1_DH_) shown in cartoon representation in same orientations as in Fig. 1C.

### Overall architecture of the PP2A(B56)-Sgo1-CPC complex

The structure of the PP2A(B56)-Sgo1-CPC complex was then characterized using cryo-electron microscopy (cryo-EM) (Fig. 1, C and D and fig. S1). The structure showed the canonical folding for the PP2A(B56) holocomplex (resolved at resolutions ranging from 2.5 to 3.5 Å), with the A subunit forming the characteristic horseshoe-shaped HEAT-repeat-containing scaffold that accommodates the catalytic C subunit and the HEAT-repeat-containing B56 regulatory subunit. The structure also showed the Sgo1_DH_ (containing Sgo1_51-106_, average resolution of 3.0 Å) engaging both the B56 and C subunits of PP2A, in agreement with the previously reported crystal structure of the PP2A(B56)-Sgo1_51-96_ complex (*42*) (Fig. 1, C and D). Compared to the crystal structure, the cryo-EM structure shows that Sgo1DH is 10 amino acids longer at the C-terminus (53–106), forming three additional helical turns (Fig. S2A).

In addition, the structure revealed an unaccounted density occupying the LxxIxE-binding pocket of PP2A(B56), with an average local resolution of 2.5 Å (Fig. 1, C and D and fig. S1E). This density extends beyond the LxxIxE-binding pocket, occupying a previously uncharacterized interaction interface between HEAT repeats 1-5 in the A subunit and HEAT repeats 4-7 in the B56 subunit (Fig. 1, C and D and fig. S1, ‘Borealin loop’ map). PP2A(B56) recognizes its binding partners through LxxIxE motifs, which engage a conserved surface groove on the B56 regulatory subunit to mediate substrate recruitment and positioning (*15, 17*). Our previously published work suggested that Sgo1 might engage the B56 LxxIxE pocket (*16*), raising the possibility that this density could be from Sgo1.

The structure also showed a poorly resolved but strong density above Sgo1_DH,_ extending up to the N-terminal region of B56 and beyond the C-terminal end of Sgo1_DH_ (Fig. 1, C and D, colored in light grey). Given that CPC can bind Sgo1 via the interaction between the Sgo1 N-terminal tail and the Survivin BIR domain (*27, 33*), and that the Sgo1 construct used for reconstitution includes an intrinsically disordered region beyond the Sgo1_DH_ (Sgo1_IDR_, comprising residues 107-155), we hypothesized that both Sgo1-tethered CPC and the Sgo1_IDR_ contribute to the heterogeneous density.

To unambiguously assign the unaccounted densities, we reconstituted the PP2A(B56)-Sgo1 complex using recombinant Sgo1_1-155_ and the PP2A(B56) holoenzyme (Fig. 1B) and determined its structure using cryo-EM (Fig. 2A). Consistent with our PP2A(B56)-Sgo1-CPC cryo-EM structure, the PP2A(B56)-Sgo1 structure revealed a well-resolved Sgo1_DH_ bridging the B56 and C subunits of PP2A (resolved at a resolution ranging from 2.5 to 3.5 Å) and a poorly resolved ‘arch-like’ density with an average resolution of 8 Å above the Sgo1_DH_ that extends beyond the C-terminal end of the Sgo1_DH_ (Fig. 2A, fig S2B and fig. S3). XL-MS data suggest that the N-terminal helices of Sgo1 (Sgo1 Nt helices) fold back and interact with the Sgo1_DH,_ explaining the ‘arch-like’ density above Sgo1_DH_ (Fig. 2B), also supported by MD simulations, (Fig. S2, C and D). Focused classification analysis revealed that the density at the C-terminus of the Sgo1_DH_ represents the averaging of multiple Sgo1_IDR_ conformations against the B56 subunit (Fig. S2E).

**Figure 2:**
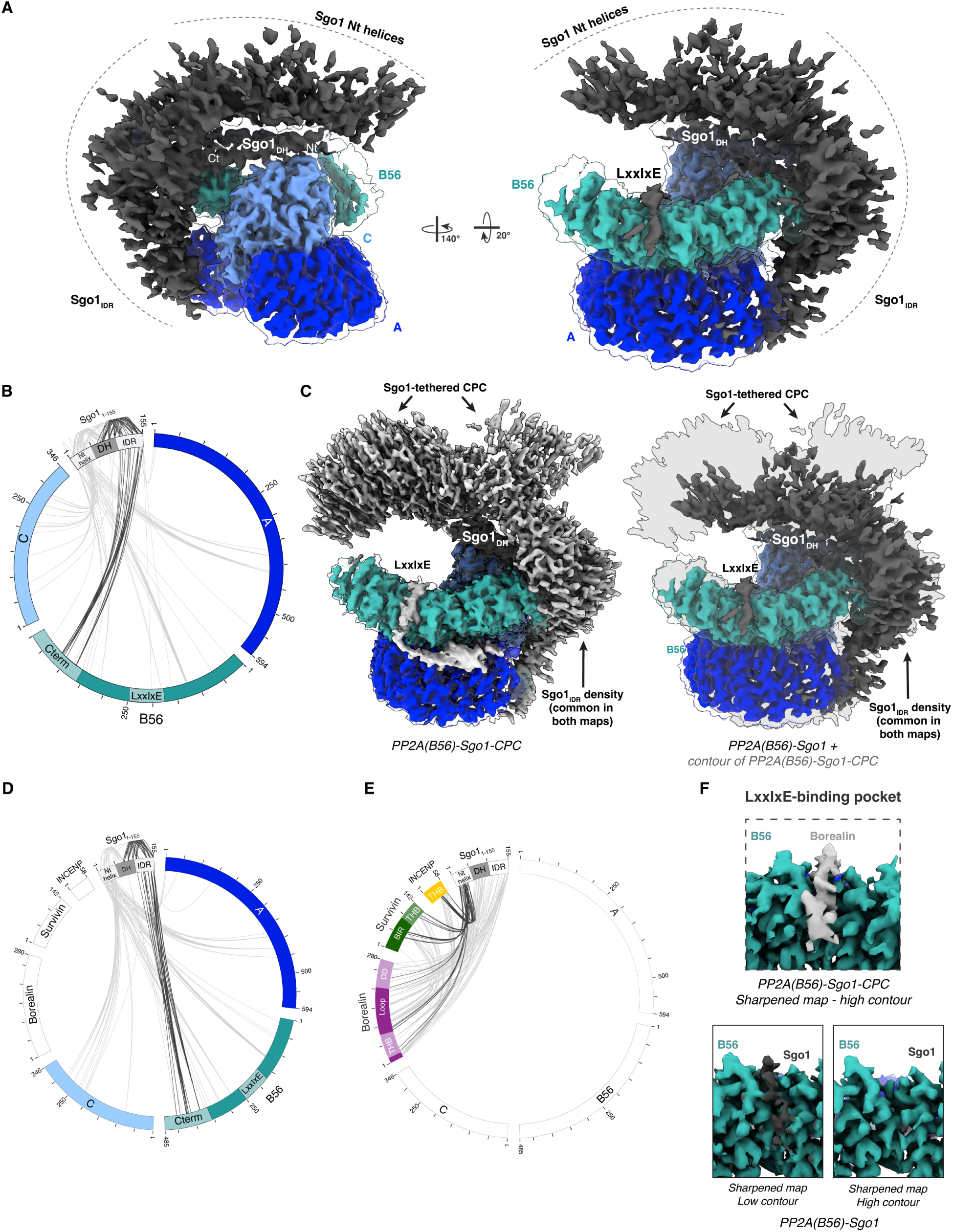
PP2A(B56)-Sgo1 structure reveals Sgo1_IDR_ and Sgo1 LxxIxE-like motif interactions with B56. **(A)** Overlay of the ‘consensus’ and ‘high-resolution’ cryo-EM maps of the PP2A(B56)-Sgo1 complex (Fig. S3). Conformationally heterogeneous densities corresponding to Sgo1_IDR_ and Nt helices are shown in dark grey and labelled. **(B)** XL-MS circular diagram of the PP2A(B56)-Sgo1 complex showing the crosslinks between Sgo1 and PP2A(B56). Crosslinks involving Sgo1_IDR_ and Sgo1_DH_ and the C-terminal region of B56 are highlighted in dark grey. **(C)** Comparison between the consensus maps of the PP2A(B56)-Sgo1-CPC_I1-58SB_ (left) and PP2A(B56)-Sgo1 (right) complexes. The silhouette of the PP2A(B56)-Sgo1-CPC_I1-58SB_ is overlaid as a transparent grey surface onto the PP2A(B56)-Sgo1 map to highlight additional CPC-derived density. **(D)** XL-MS circular diagram of the PP2A(B56)-Sgo1-CPC_I1-58SB_ complex showing the crosslinks between Sgo1 and PP2A(B56). Crosslinks involving Sgo1_IDR_ are shown in dark grey. **(E)** XL-MS circular diagram of the PP2A(B56)-Sgo1-CPC_I1-58SB_ complex showing the crosslinks between Sgo1 and CPC_I1-58SB_. Crosslinks involving Sgo1 N-terminal helices are shown in dark grey. **(F)** Close-up views of the LxxIxE-binding pocket in the PP2A(B56)-Sgo1-CPC_I1-58SB_ structure (top) and PP2A(B56)-Sgo1 structure (bottom). Two different contours are shown for the PP2A(B56)-Sgo1 map.

In addition, XL-MS analysis and MD simulations show that Sgo1_IDR_ folds back and establishes intra-molecular contacts with itself and with Sgo1_DH_ (Fig. 2B and fig. S2, C and D).

An overlay of the density maps corresponding to PP2A(B56)-Sgo1-CPC and PP2A(B56)-Sgo1 shows that, in the PP2A(B56)-Sgo1-CPC complex, the density above Sgo1_DH_ occupies a much larger conformational space (Fig. 2C). The heterogeneous density corresponding to the Sgo1_IDR_ is present in both structures (also consistent with XL-MS analysis, fig. 2D), suggesting that the additional heterogeneous density observed for the PP2A(B56)-Sgo1-CPC complex corresponds to two CPC triple-helical bundles (CPC_THB_, fig. 1A, composed of INCENP_1-58_, Survivin_80-142_ and Borealin15-76) tethered to the N-terminal tails of the Sgo1 dimer, hovering above the Sgo1_DH_. Consistent with this, XL-MS analysis showed prominent contacts between the N-terminal helices of Sgo1 and the CPC_THB_ (Fig. 2E).

The PP2A(B56)-Sgo1 structure also showed density at the LxxIxE-binding pocket of B56, likely corresponding to Sgo1, visible only at lower contour levels (Fig. 2F and fig. S2F). Notably, in this structure, the density observed at the A-B56 interface of the ternary complex was absent (Fig. 2C). AlphaFold 3 (AF3) model prediction suggested that Sgo1_141-145_, containing the ‘LxxIxE’-like motif I_141_-x-L-x-E_145_, might engage the pocket (Fig. S2G). Thus, while this agrees with our previous study that Sgo1 might engage the LxxIxE-binding pocket, the interaction is likely to be weak or transient. In contrast, the unaccounted density at the LxxIxE-binding pocket in the PP2A(B56)-Sgo1-CPC is well resolved (Fig. 2F), and it extends beyond the LxxIxE-binding pocket at the interface between A and B56 subunits (Fig. 1, C and D and fig. 2C). This led us to hypothesize that a previously uncharacterized ‘LxxIxE’-containing region of the CPC directly interacts with PP2A(B56).

### CPC engages PP2A(B56) via an ‘LxxIxE-like’ motif present in Borealin

XL-MS analysis of the PP2A(B56)-Sgo1-CPC complex identified crosslinks consistent with the density in the LxxIxE-binding pocket and beyond originating from the Borealin subunit of CPC (Fig. 3A). Sequence analysis identified two candidate LxxIxE-like motifs within the Borealin loop region that are highly conserved in mammals: V_116_-x-x-V-x-E_121_ and D_120_-x-x-I-x-E_125_ (Fig. 3B). To identify which candidate motif occupies the LxxIxE-binding pocket of PP2A(B56), we assessed the fitting of the two motifs into the cryo-EM density, with both motifs showing strong density correlation (Fig. S4A). Nevertheless, unlike the VxxVxE motif, the DxxIxE motif contains a downstream stretch of negatively charged residues (E_126_-E_127_-E_128_ downstream DxxIxE vs. M_122_-I_123_-V_124_ downstream VxxVxE) that would establish favorable interactions with the cluster of B56 basic residues downstream of the pocket (Fig. 3, B and C and fig. S4B). XL-MS analysis coupled to AlphaFold3 modelling (AF3x, (*43*)) enabled the assignment of residues downstream of the Borealin DxxIxE motif (Borealin_135-145_) binding along the A-B56 interface with good density correlation for the Cα backbone of the Borealin loop (Fig. 3D).

**Figure 3:**
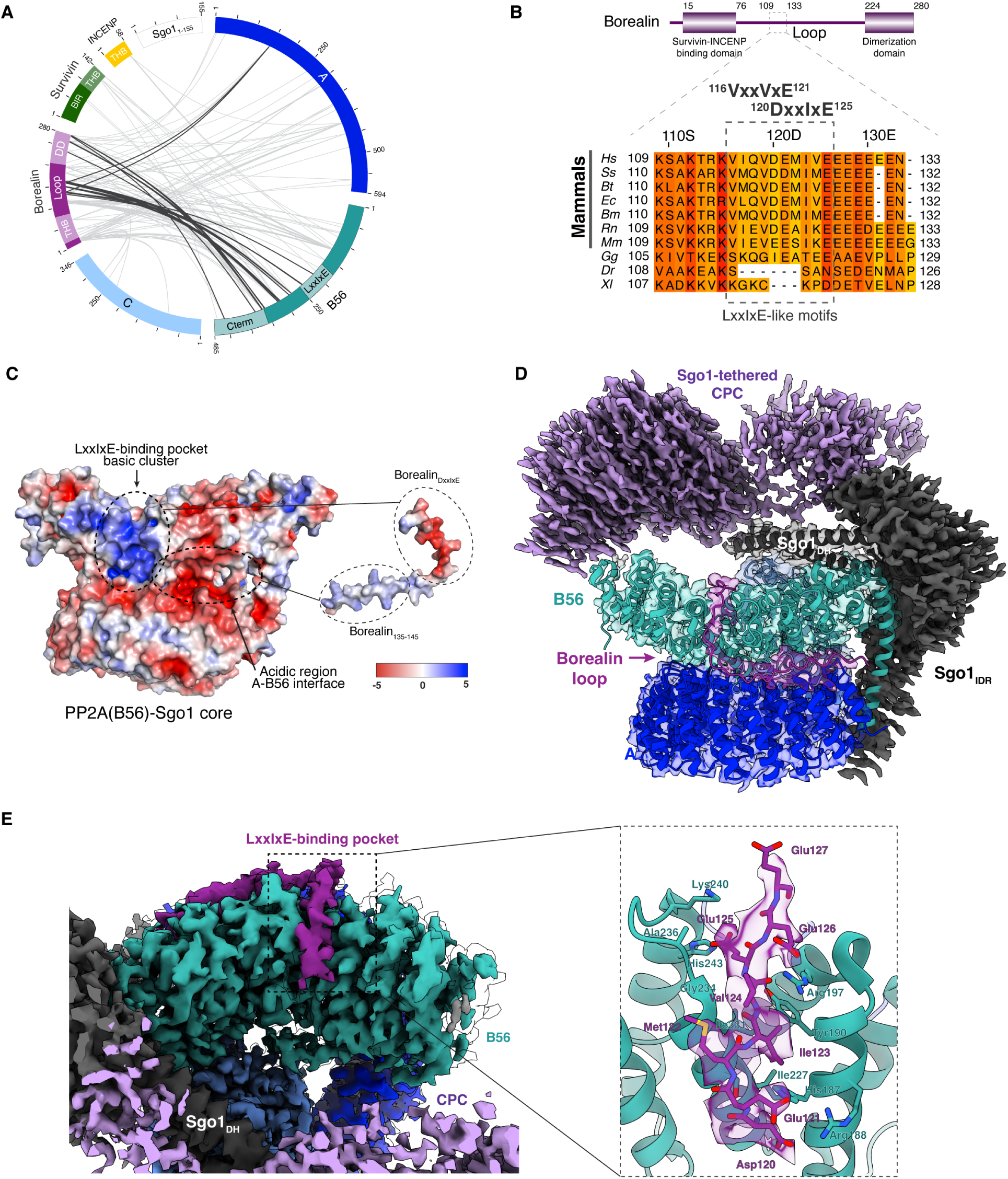
Borealin contains an ‘LxxIxE’-like motif that interacts with B56 and a downstream loop that interacts at a novel A-B56 interface. **(A)** XL-MS circular diagram of the PP2A(B56)-Sgo1-CPC_I1-58SB_ complex showing the crosslinks between PP2A(B56) and CPC_I1-58SB_. Crosslinks involving the Borealin loop and dimerization domain (DD) are shown in dark grey. **(B)** Amino acid conservation of Borealin_109-133_ containing LxxIxE-like motifs (well conserved - darker orange and less conserved - yellow): Homo sapiens (Hs), Sus scrofa (Ss), Bos taurus (Bt), Equus caballus (Ec), Balaenoptera musculus (Bm), Rattus norvegicus (Rn), Mus musculus (Mm), Gallus gallus (Gg), Danio rerio (Dr) and X. laevis (Xl). **(C)** Adaptive Poisson-Boltzmann Solver (APBS) surface electrostatic potential and corresponding interacting regions of the PP2A(B56)-Sgo1 (left) and the Borealin loop containing the DxxIxE motif (right). **(D)** Overlay of the ‘consensus’, ‘high-resolution’ and ‘Borealin loop’ cryo-EM maps of the PP2A(B56)-Sgo1-CPC_I1-58SB_ complex (Fig. S1). Conformationally heterogeneous densities corresponding to Sgo1_IDR_ and Sgo1-tethered CPC, are shown in dark grey and lilac, respectively. The Borealin densities at the LxxIxE-binding pocket and on the A-B56 interface are shown in purple. The AF3x model is shown in cartoon representation within the corresponding regions of the map shown with transparency. **(E)** Top view of the overlay of ‘consensus’, ‘high-resolutio’n and ‘Borealin loop’ maps of the PP2A(B56)-Sgo1-CPC_I1-58SB_ complex (left) and close-up view of the inter-molecular contacts between Borealin DxxIxE and the B56 LxxIxE-binding pocket with corresponding cryo-EM density (left).

Eight Borealin residues (Asp_120_-Glu_127_) could be confidently assigned within the map, of which Asp_120_, Ile_123_, Val_124_ and Glu_125_ establish backbone-and side chain-mediated interactions with the B56 LxxIxE-binding pocket residues His_187_, Arg_188_, Tyr_190_, Arg_197_, Ile_227_, Ile_231_, Gly_234_, Ala_236_, Lys_240_, His_243_ (Fig. 3E). Unlike canonical LxxIxE motifs, the first residue is an aspartate rather than a leucine. In our structure, Borealin Asp_120_ forms hydrogen bonding interactions with B56 His_187_ and Arg_188_, thereby defining a previously uncharacterized variant of the LxxIxE motif. As mentioned earlier, Borealin_126-132_ packs against the basic cluster downstream of the LxxIxE-binding pocket. In addition to this, a positively charged stretch of Borealin residues 135-145 engages an acidic region of the A-B56 interface (Fig. 3C).

The observation that Borealin establishes prominent contacts with the LxxIxE-binding pocket of PP2A(B56) and beyond raised the possibility of a direct interaction between PP2A(B56) and CPC.

### Independent of Sgo1, PP2A(B56) and CPC directly interact through Borealin ‘DxxIxE’ and the dimerization domain

To test whether CPC and PP2A(B56) complexes can interact directly, we performed SEC analysis with recombinant PP2A(B56) holocomplex and CPC_I1-58SB_. SEC analysis showed that, independent of Sgo1, PP2A(B56) and CPC form a complex that eluted at 11.5 ml compared to PP2A(B56) and CPC alone, which eluted at 13.0 ml and 11.8 ml, respectively (Fig. 4A). To understand the structural basis for this interaction, we determined the cryo-EM structure of the PP2A(B56)-CPC complex (resolution ranging from 2.5 to 5 Å). The cryo-EM structure revealed a well-defined density at the LxxIxE-binding pocket and along the interface between PP2A A and B56 subunits identical to those observed in the PP2A(B56)-Sgo1-CPC complex structure, unambiguously confirming the engagement of the Borealin loop (Fig. 4, B and C and fig. S5, A and D). Consistent with this, a CPC construct lacking the N-terminal fragment of the Borealin loop (CPC_I1-58SBΔ110-188_) failed to interact with PP2A(B56) by SEC (Fig. S6A).

**Figure 4:**
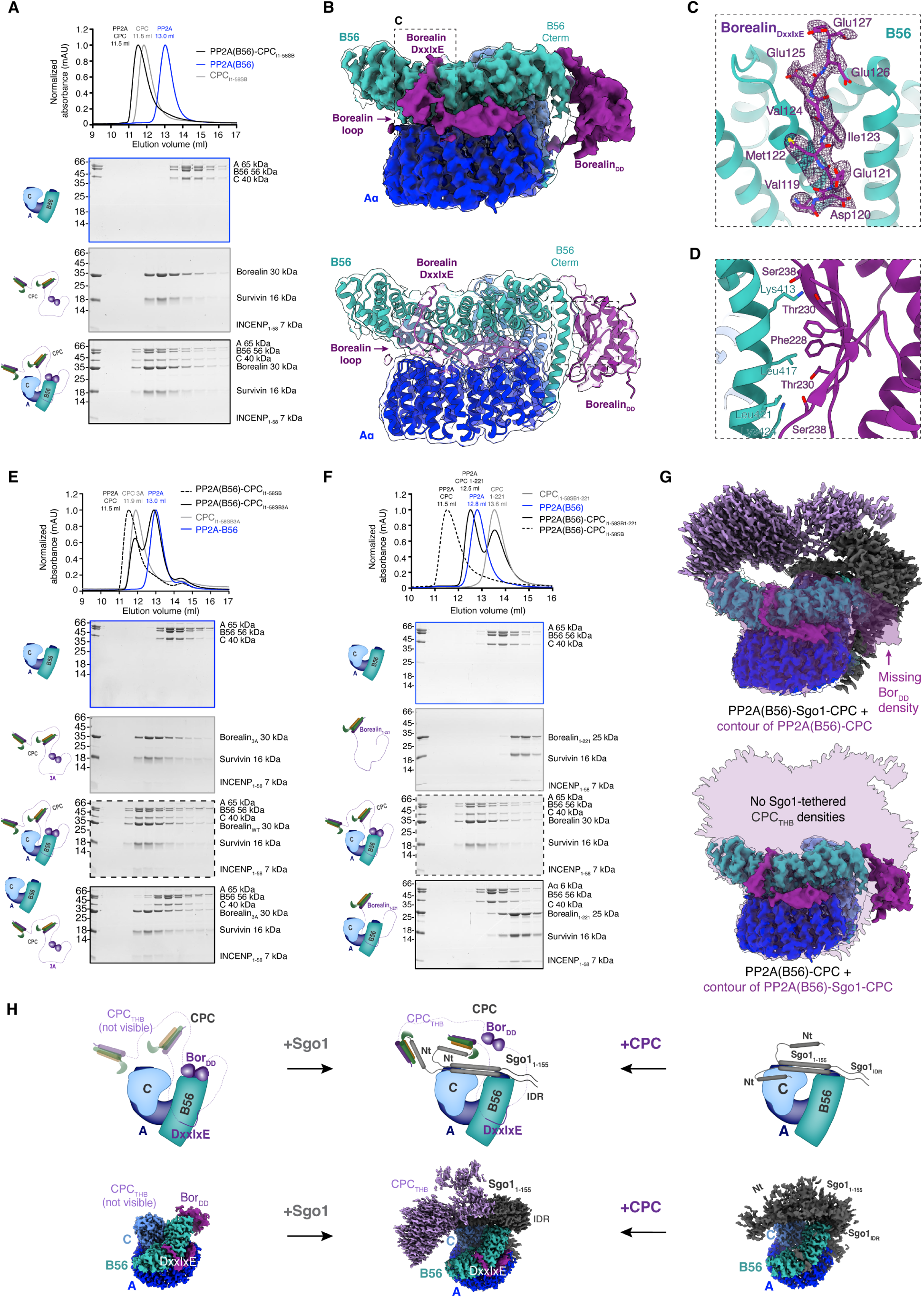
Structural basis for Sgo1 independent PP2A(B56)-CPC assembly. **(A)** SEC profiles and corresponding SDS-PAGEs of PP2A(B56) (blue), CPC_I1-58SB_ (grey) and PP2A(B56)-CPC_I1-58SB_ (black). **(B)** Overlay of the ‘high-resolution’ and the ‘Borealin loop and dimerization domain’ maps of the PP2A(B56)-CPC_I1-58SB_ complex (top, Fig. S6) with corresponding model fitted and maps shown with transparency (bottom). **(C)** Inter-molecular contacts between Borealin DxxIxE and the LxxIxE-binding pocket of B56, with corresponding cryo-EM density for Borealin shown as a mesh. **(D)** Inter-molecular contacts between Borealin_DD_ and the C-terminal region of B56. **(E)** SEC profiles and corresponding SDS-PAGEs of PP2A(B56) (blue), CPC_I1-58SB3A_ (grey), the PP2A(B56)-CPC_I1-58SB_ complex (dashed black) and PP2A(B56)-CPC_I1-58SB3A_ (solid black). **(F)** SEC profiles, and corresponding SDS-PAGEs of PP2A(B56) (blue), CPC_I1-58SB1-221_ (grey), PP2A(B56)-CPC_I1-58SB_ (dashed black) and PP2A(B56)-CPC_I1-58SB1-221_ (solid black). The chromatogram and SDS-PAGEs corresponding to the PP2A(B56)-CPC_I1-58SB_ complex from fig. 4A are used in this panel for direct comparison. **(G)** Comparison between the consensus maps of the PP2A(B56)-Sgo1-CPC_I1-58SB_ (top) and PP2A(B56)-CPC_I1-58SB_ (bottom) complexes. Transparent purple silhouettes of the PP2A(B56)-CPC_I1-58SB_ (top) and PP2A(B56)-Sgo1-CPC_I1-58SB_ (bottom) complexes are overlayed. **(H)** Representative cartoons for the PP2A(B56)-CPC_I1-58SB_ (left), PP2A(B56)-Sgo1-CPC_I1-58SB_ (middle) and PP2A(B56)-Sgo1 (right) complexes (top) and corresponding cryo-EM structures (bottom).

To further confirm the role of the Borealin DxxIxE motif in stabilizing the assembly of the PP2A(B56)-CPC complex, we mutated Asp_120_, Ile_123_ and Glu_125_ of Borealin to Ala (Borealin_D120A/I123A/E125A_, Borealin_3A_) and tested complex formation by SEC. CPC_I1-58SB_ containing the Borealin_3A_ mutant (CPC_I1-58SB3A_) abolished binding to PP2A(B56) (Fig. 4E), whereas mutation of the alternative LxxIxE-motif VxxVxE (CPC_I1-58SB_ containing Borealin_V116A/V119A/E121A_) still maintained the ability to interact with PP2A(B56) (Fig. S6B). Likewise, the previously characterized B56 R197E LxxIxE-binding pocket mutation (*16*), which disrupts recognition of LxxIxE-containing binding partners, completely abolished CPC binding in the context of the B56 subunit alone (Fig. S6C). Together, these findings demonstrate that the binding of Borealin DxxIxE motif to the PP2A LxxIxE-binding pocket is crucial for PP2A(B56)-CPC interaction.

The structure also revealed an unaccounted density next to the C-terminal region of B56 (Fig 4B), previously reported to play a role in the isoform-specific localization of PP2A(B56) during mitosis (*19*). XL-MS data obtained for the PP2A(B56)-CPC complex revealed extensive contacts between the Borealin dimerization domain (Borealin_DD_) and the C-terminal region of B56 (Fig. S6D). In agreement with this, the overall dimensions of the unaccounted density matched those of Borealin_DD,_ which could be placed within this density using rigid-body fitting (Fig. 4B and fig. S5E). In this mode of binding, a hydrophobic core composed of Borealin Phe_228_ flanked by polar interactions involving Borealin Thr_230_ and Ser_238_ would mediate the interaction with the C-terminal helix of B56 (Fig. 4D). Together, these findings indicate that Borealin dimerization domain provides an additional interaction interface stabilizing the PP2A(B56)-CPC complex. To test this hypothesis, we performed SEC experiments with a CPC construct lacking the Borealin_DD_ (CPC_I1-58SB1-221_). Removal of the Borealin_DD_ markedly reduced complex formation in the context of the PP2A(B56)-CPC complex, as evidenced by the absence of a pronounced shift in the SEC elution profile compared to CPC containing full-length Borealin (Fig. 4F).

Comparison of the PP2A(B56)-Sgo1-CPC and PP2A(B56)-CPC structures revealed that, in the latter complex, the Borealin_DD_ density is replaced by the density corresponding to the Sgo1_IDR_ (Fig. 4G), indicating that in the presence of Sgo1, the Borealin_DD_ cannot interact with the C-terminal helix of the B56 subunit. Moreover, poorly resolved densities observed above Sgo1_DH_ in the PP2A(B56)-Sgo1-CPC complex are absent in the PP2A(B56)-CPC structure. This further confirms that the poorly resolved density observed in the PP2A(B56)-Sgo1-CPC complex corresponds to the CPC_THB_ tethered to the N-terminal tail of Sgo1, an observation further supported by MD simulations of the PP2A(B56)-Sgo1-CPC complex (Fig. S7, A and B). Therefore, Sgo1-mediated tethering of the CPC_THB_ would provide additional stabilization in the absence of Borealin_DD_ interaction with the C-terminal region of B56.

Altogether, the integrative analysis of our structural and biochemical data for the PP2A(B56)-Sgo1, PP2A(B56)-CPC, and PP2A(B56)-Sgo1-CPC complexes allowed us to map the PP2A(B56)-CPC interactions within the ternary complex, and to define the role of Sgo1 in its assembly (Fig. 4H). Our observations suggest that the CPC engages PP2A(B56) through two anchoring points. One of them is the interaction between the Borealin DxxIxE and the PP2A(B56) LxxIxE-binding pocket, which is observed both in the presence and absence of Sgo1. Unlike the first anchoring point, the second one is variable and depends on the presence of Sgo1 (PP2A(B56)-CPC vs PP2A(B56)-Sgo1-CPC). In the absence of Sgo1, the interaction between the Borealin_DD_ and the C-terminal region of B56 would constitute the second anchoring point. This is not compatible with the presence of Sgo1, as the Sgo1_IDR_ displaces the Borealin dimerization domain from its interaction with the C-terminal region of the B56 subunit. This is compensated by a new anchoring point that tethers CPC to the N-terminal tail of Sgo1 via its interaction with the BIR domain of Survivin. This hypothesis is consistent with the loss of CPC binding in the context of the Sgo1_Nmut_, where the second anchoring point involving the N-terminal tail of Sgo1 and Survivin is disrupted (Fig. S7C).

### Perturbing PP2A(B56)-CPC interaction compromises chromosome segregation fidelity

We first assessed the interaction between PP2A(B56) and CPC in cells by performing GFP-Trap immunoprecipitation assays using a doxycycline-inducible HeLa cell line expressing YFP-PP2A(B56) and transiently expressing either Myc-Borealin_WT_ or the PP2A-binding mutant Myc-Borealin_D120A/I123A/E125A_ (Myc-Borealin_3A_). Myc-Borealin_WT_ efficiently co-immunoprecipitated with YFP-PP2A(B56), whereas the Borealin 3A mutant exhibited markedly reduced binding, confirming that the Borealin DxxIxE motif mediates the PP2A(B56)-CPC interaction in cells (Fig. 5A).

**Figure 5:**
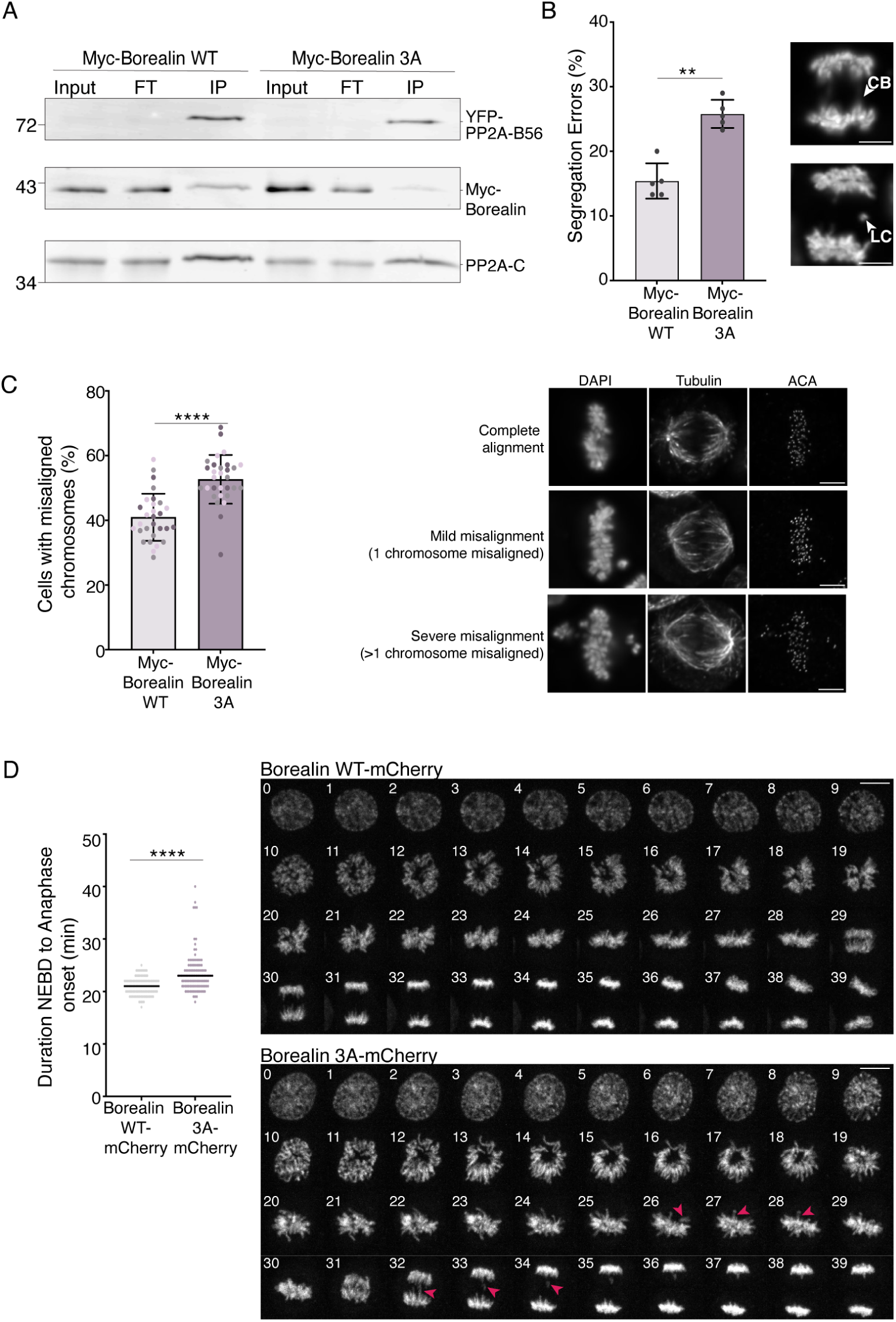
CPC–PP2A(B56) interaction is essential for faithful chromosome segregation. **(A)** GFP-Trap immunoprecipitation of YFP-PP2A(B56) from HeLa cells, assessing its interaction with Myc-tagged Borealin WT (Myc-Borealin_WT_) and 3A mutant (Myc-Borealin_D120A/I123A/E125A_). The PP2A catalytic subunit (PP2A-C) was used as a control. **(B)** Quantification of anaphase chromosome bridges (CB) or lagging chromosomes (LC) in HeLa Kyoto cells following Borealin depletion and subsequent expression of either Myc-tagged Borealin WT or Borealin 3A. DNA was visualised using DAPI. Scale bar, 5 µm. Data represent the mean ± SD from five independent experiments, at least 100 anaphases were analysed per replicate, Mann-Whitney U test; **, P ≤ 0.01. **(C)** Quantification of cells with misaligned chromosomes in HeLa Kyoto cells subjected to a chromosome biorientation assay. DNA was visualised using DAPI. Scale bar, 5 µm. Data represent the mean ± SD from three independent experiments, with n ≥ 155 of metaphases analysed per replicate, Mann-Whitney U test; ****, P ≤ 0.0001. **(D)** Quantification and representative images (1 min intervals) from live-cell imaging showing mitotic progression in hTERT-RPE1 cells expressing Borealin WT-mCherry (light pink) or Borealin 3A-mCherry (purple). In representative images, 0 min was defined as the onset of visible chromosome condensation. Mitotic timing was quantified in at least 110 cells per condition across three independent biological replicates. Scatter plots show the time elapsed (in minutes) from nuclear envelope breakdown (NEBD) to anaphase onset. Each dot represents an individual cell, and horizontal black lines indicate median values. Mann-Whitney U test; ****, P ≤ 0.0001. Red arrows highlight congression defects and segregation errors.

To evaluate the requirement of the PP2A(B56)-CPC interaction in ensuring accurate chromosome segregation, we performed small interfering RNA (siRNA)-rescue assays in HeLa Kyoto and replacement assays in hTERT-RPE1 cells using the Borealin_3A_ mutant. In HeLa Kyoto cells, endogenous Borealin was depleted by siRNA, followed by transient expression of either Myc-tagged Borealin_WT_ or Myc-tagged Borealin_3A_ (Fig. S8A). In hTERT-RPE1 cells, cell lines expressing Borealin-mClover in which endogenous Borealin could be replaced with either Borealin_WT_-mCherry or Borealin_3A_-mCherry were generated (Fig. S8B).

Perturbation of the PP2A(B56)-CPC interaction, although it did not affect bulk CPC levels at centromeres (Fig. S8, C and D), resulted in mitotic defects in both cell lines. Compared with Borealin_WT_-expressing cells, Borealin_3A_ cells displayed significantly increased chromosome segregation errors during anaphase, including chromosome bridges and lagging chromosomes (Fig. 5B and fig. S8E). To assess whether these defects arose from impaired chromosome biorientation, HeLa and RPE1 cells were released from a monastrol-induced mitotic arrest and chromosome alignment was quantified. Borealin3A expression significantly increased the frequency of chromosome congression defects relative to Myc-Borealin_WT_ (Fig. 5C and fig. S8F). Consistent with these observations, live-cell imaging revealed a tendency to prolong the interval between NEBD and anaphase onset in Borealin_3A_-expressing cells (Fig. 5D and fig. S8G), indicative of delayed mitotic progression.

Together, these findings demonstrate that the direct interaction between PP2A(B56) and CPC is required to ensure accurate mitosis. Disruption of this interaction compromises chromosome congression, delays mitotic progression and ultimately reduces the fidelity of chromosome segregation.

## Discussion

Precise spatiotemporal regulation of reversible phosphorylation orchestrates each stage of mitosis, including maintenance of sister chromatid cohesion, chromosome biorientation through the stabilization of correct chromosome-microtubule attachments, and timely sister chromatid separation. Centromeres and kinetochores act as signaling hubs where the coordinated activities of kinases, including Aurora B (the enzymatic core of the CPC), CDK1, PLK1 and Mps1, and phosphatases like PP2A and PP1, dynamically regulate phosphorylation of centromere and kinetochore substrates, such as histones and the KMN network (NDC80/HEC1, DSN1, KNL1). This balance must be robust enough to eliminate erroneous attachments while remaining sufficiently responsive to stabilise correct attachments and permit anaphase onset. For example, phosphorylation of NDC80 promotes the destabilisation of inappropriate attachments and contributes to checkpoint signalling, whereas its timely dephosphorylation supports attachment stabilisation, chromosome alignment and mitotic progression.

The Aurora B-containing CPC and PP2A(B56) constitute a major pair of opposing enzymatic activities that regulate kinetochore-microtubule attachment stability and chromosome segregation (*37, 44, 45*). Sgo1 interacts independently with both PP2A(B56) and CPC and is therefore well positioned to coordinate their recruitment and activity at centromeres and kinetochores. However, whether Sgo1 acts as a scaffold that simultaneously engages these opposing enzymes and how this arrangement regulates their local balance remained unclear.

Here, we show that PP2A(B56) and CPC form an integrated dynamic signaling platform. We identify two interfaces that mediate PP2A(B56)-CPC association. The primary interface is formed by a previously uncharacterized DxxIxE motif in Borealin, which engages the canonical LxxIxE-binding pocket of B56 and provides the principal anchoring site. A second interface between the Borealin_DD_ and the C-terminal region of B56 further stabilizes the complex. Together, these contacts establish a bipartite interaction that directly couples PP2A(B56) to CPC.

Our data further suggest that Sgo1 remodels, rather than simply promotes, the PP2A(B56)-CPC interaction. In the presence of Sgo1, the C-terminal IDR of Sgo1 interferes with the Borealin_DD_-B56 interaction while maintaining the Borealin DxxIxE-B56 contact. The resulting loss of Borealin_DD_-B56 interaction appears to be compensated by the interaction of Sgo1 N-terminal tail with Survivin BIR domain. This arrangement provides a mechanistic explanation for how Sgo1 can coordinate PP2A(B56) and CPC, two opposing enzymatic activities, without disrupting their association, while simultaneously altering the architecture and likely the functionality of the resulting complex.

The integration of PP2A(B56) and CPC within a single assembly has several important implications. First, their direct association could enable reciprocal recruitment or retention at centromeres and kinetochores, thereby influencing the local abundance, residence time and relative positioning of the two activities. Their proximity could also facilitate phosphoregulation: PP2A(B56) could dephosphorylate CPC components, whereas Aurora B could phosphorylate subunits of the PP2A(B56) holoenzyme, potentially altering activity, localization, or interactions of either complex. Such cross-regulation could establish local feedback mechanisms that tune their functional balance, extending their relationship beyond opposing effects on shared substrates. Sgo1 may therefore coordinate these activities not simply by recruiting each complex independently, but by assembling and remodeling a shared signaling module that influences both their access to substrates and their regulation of one another.

Second, CPC binding may influence substrate access to PP2A(B56). Borealin engages the conserved LxxIxE-binding pocket, which also mediates recognition of key regulators, including BubR1, RepoMan and Kif4A (*15–18, 39, 46*). Occupancy of this pocket could therefore compete with other LxxIxE-containing proteins and influence substrate recruitment, exchange and dephosphorylation kinetics. As LxxIxE-mediated interactions contribute to both substrate recognition and phosphatase targeting, CPC binding could couple the localization of PP2A(B56) to changes in the repertoire of substrates accessible to it (*15, 19*).

An additional potential layer of regulation is phosphorylation-dependent control of PP2A(B56)-CPC association itself. Phosphorylation within or adjacent to LxxIxE motifs can enhance B56 binding, providing an established mechanism for coupling kinase activity to phosphatase recruitment (*15, 17*). This raises the possibility that phosphorylation at or near the Borealin DxxIxE motif, or at the secondary interface, could modulate the affinity and organization of the PP2A(B56)-CPC assembly. Together with potential reciprocal phosphoregulation within the complex, such a mechanism could couple assembly and remodeling to changes in the functional state of its constituent enzymes. Identifying the relevant phosphorylation sites and determining their effects on complex formation and activity will be important for establishing how this module is regulated during mitosis.

Overall, our findings suggest that mitotic kinase-phosphatase balance is governed not only by the independent recruitment of opposing enzymes, but also by their physical organization within shared regulatory assemblies. The structural framework presented here enables direct tests of how assembly, remodeling and reciprocal phosphoregulation shape the spatiotemporal control of mitotic substrate phosphorylation to promote accurate chromosome segregation.

## Materials and Methods

### Protein expression and purification of PP2A(B56γ)

PP2A(B56γ) was expressed and purified as described in Vit et al., 2022 (*47*).

In brief, GST-tagged Aα and B56γ subunits were expressed and purified separately in the same conditions. Aα and B56γ in pGEX-6P-1 vectors were expressed in *E. coli* BL21 Rosetta cells grown in Terrific Broth (TB) media at 37 °C until OD600 reached ∼1.0 and induced overnight at 18 °C by the addition of 0.35 mM Isopropyl-β-D-1-thiogalactopyranoside (IPTG). Cultures were harvested by centrifugation at 4000 g for 10 min at 4 °C and resuspended in lysis buffer (20 mM Tris-HCl pH 8.0, 500 mM NaCl, 2 mM dithiothreitol [DTT]) supplemented with 10 μg/ml DNase and 1 mM phenylmethylsulphonyl fluoride (PMSF). After cell lysis by sonication, the lysate was clarified by centrifugation at 22,000 rpm for 50 min and incubated for 2 h at 4 °C with 20 ml of GSH slurry (Cytiva) pre-equilibrated with lysis buffer. The protein-bound GSH resin was then washed with lysis buffer and high salt buffer (containing 20 mM Tris-HCl pH 8.0, 1000 mM NaCl, 50 mM KCl, 10 mM MgCl2, 2 mM ATP, 2 mM DTT). A beads-bound elution was performed overnight by the addition of 2 mg of GST-tagged 3C protease and eluted protein was concentrated using an Amicon Ultra 15 10 kDa MWCO centrifugal unit (Millipore). Protein was applied onto a HiLoad Superdex 200 16/600 pg column (Cytiva) pre-equilibrated with 20 mM Tris-HCl pH 8.0, 500 mM NaCl, 5 % (V/V) glycerol, 2 mM DTT. Fractions containing pure protein were analyzed by SDS-PAGE, pooled and snap frozen with liquid nitrogen.

The MultiBac system (*48*) was used to generate His-tagged Cα D88N subunit. For this, high titer baculoviruses (V2) were generated using Sf9 cells and used to infect Hi5 cells grown at a density of 2×10^6^ cells/ml. Cells were infected for 48 h with V2 at a 1:25 ratio and harvested by centrifugation at 4000 g for 20 min at 4 °C. Insect cell pellets were resuspended in lysis buffer containing 20 mM Tris-HCl pH 8.0, 250 mM NaCl, 35 mM imidazole and 2 mM β-mercaptoethanol (β-ME) and disrupted by sonication. After clarifying the lysate by centrifugation, the soluble overexpressed protein was loaded for 3 h at 4 °C onto 10-ml Ni2+-NTA slurry (Cytiva) previously equilibrated with lysis buffer. Beads were subsequently washed with lysis buffer and high salt buffer (20 mM Tris-HCl pH 8.0, 1 M NaCl, 35 mM imidazole, 50 mM KCl, 10 mM MgCl2, 2 mM ATP, 2 mM β-ME) before elution with buffer containing 20 mM Tris-HCl pH 8.0, 250 mM NaCl, 400 mM imidazole and 2 mM β-ME.

Pure Aα and B56γ subunits and affinity-purified Cα subunit were mixed at a 1:1:1 molar ratio and dialyzed overnight at 4 °C against buffer containing 20 mM Tris-HCl pH 8.0, 150 mM NaCl, 5 % (V/V) glycerol and 2 mM DTT. To remove contaminants and the excess of B56γ subunit, the dialysed ample was applied onto a 5-ml HiTrap Q HP anion exchange chromatography column (Cytiva) and eluted using a gradient of high salt buffer (20 mM Tris-HCl pH 8.0, 1000 mM NaCl, 5 % (V/V) glycerol, 2 mM DTT). After ion exchange, the PP2A(B56γ) holocomplex co-eluting with excess of Aα subunit were and gel filtered using a HiLoad Superdex 200 16/600 pg column (Cytiva) in 20 mM Tris-HCl pH 8.0, 250 mM NaCl, 5 % (V/V) glycerol, 2 mM DTT buffer. Fractions containing the pure stoichiometric PP2A(B56γ) holocomplex were pooled and snap frozen in liquid nitrogen.

### Protein expression and purification of CPC_I1-58SB_ and Sgo1_1-155_

The vectors used to express the CPC were: pMCNcs-INCENP_1–58_ for untagged INCENP_1-58_, pRSET-HisGFP-3C-Survivin for full-length Survivin and pETMC-His-TEV-Borealin for full-length Borealin WT, 3A (D120A/I123A/E125A), VxxVxE 3A (V116A/V119A/E121A) and Borealin Δ110-180. The CPC was expressed and purified as described in Gireesh et al., 2025, Abad et al., 2022 and Abad et al., 2019 (*33–35*).

In brief, vectors containing INCENP, Survivin and Borealin were co-transformed into *E. coli* BL21 (DE3) pLysS and grown in TB media at 37 °C. After OD ∼ 0.8, 0.35 mM IPTG was used to induced protein expression O/N at 18 °C. Cultures were pelleted by centrifugation, resuspended in lysis buffer (25 mM Hepes pH 7.5, 500 mM NaCl, 25 mM imidazole, and 2 mM β-mercaptoethanol –supplemented with 0.01 mg/ml DNase (Sigma-Aldrich), 1 mM PMSF and complete EDTA-free cocktail tablets (Roche)) and disrupted by sonication. After lysate clarification by centrifugation, the sample was applied onto a HisTrap FF 5-ml column (Cytiva), washed with lysis buffer and high salt buffer (25 mM Hepes pH 7.5, 1 M NaCl, 25 mM imidazole, 50 mM KCl, 10 mM MgCl, 1 mM ATP, 2 mM β-ME) and was eluted using 25 mM Hepes pH 7.5, 500 mM NaCl, 400 mM imidazole, 2 mM β-ME. After overnight dialysis (25 mM Hepes pH 7.5, 250 mM NaCl, 4 mM DTT) in the presence of TEV and 3C, the HisGFP tag and degraded or non-stoichiometric versions of the complex were separated from the intact stoichiometric complex by cation exchange using a HiTrap SP HP column (Cytiva). Protein was concentrated and further purified by SEC using a HiLoad Superdex 200 pg column (Cytiva) pre-equilibrated with 25 mM Hepes pH 8.0, 250 mM NaCl, 5 % glycerol, 4 mM DTT. Pure stoichiometric CPC complexes were snap frozen.

Sgo1_1-155_ was expressed and purified as described in Abad et al. 2022. In brief, Sgo1_1-155 WT/Nmut_ were recombinantly expressed as N-terminal intein-tagged proteins (IMPACT™ system – NEB) in E. coli BL21 (DE3) cells. Cells were grown in TB media at 37 °C until OD600 reached ∼1.0 and protein expression was induced overnight at 18 °C by the addition of 0.35 mM IPTG. Cultures were harvested by centrifugation, resuspended in lysis buffer (20 mM Tris-HCl pH 7.5, 500 mM NaCl, 1 mM ethylenediaminetetraacetic acid (EDTA)) and disrupted by sonication. After clarification, the soluble fraction was incubated for 2 h at 4 °C with 60 ml of chitin beads (NEB) and subsequently washed with lysis buffer and high salt buffer (20 mM Tris-HCl pH 7.5, 1000 mM NaCl, 1 mM ATP, 1 mM EDTA). Protein was eluted overnight from the beads by inducing the self-splicing of the tag with buffer containing 20 mM Tris-HCl pH 7.5, 500 mM NaCl, 50 mM DTT at room temperature. Eluted protein was dialyzed for ∼24 hours at 4 °C (20 mM Tris-HCl pH 7.5, 125 mM NaCl, 50 mM arginine, 50 mM glutamate and 2 mM DTT), applied to a HiTrap SP HP cation exchange chromatography column (Cytiva) and eluted using a salt gradient (20 mM Tris-HCl pH 7.5, 1000 mM NaCl, 50 mM arginine, 50 mM glutamate and 2 mM DTT). The protein was concentrated and injected into a Superdex 200 Increase 10/300 column (Cytiva) pre-equilibrated with SEC buffer (25 mM Hepes pH 7.5, 250 mM NaCl, 50 mM arginine, 50 mM glutamate and 2 mM DTT). Pure protein was snap-frozen with liquid nitrogen.

### Interaction trials by SEC

To reconstitute the PP2A(B56γ)-Sgo1_1-155_ complex, 200 μg of PP2A(B56γ) were mixed with a two-molar excess of Sgo1_1-155_ and incubated for 30 min at 4 °C prior to injection in a Superdex 200 Increase 10/300 column pre-equilibrated with 25 mM Hepes pH 8.0, 150 mM NaCl, 5 % (V/V) glycerol and 2 mM DTT. For the PP2A(B56γ)-Sgo1_1-155_-CPC_I1-58SB_ complex, fractions containing the PP2A(B56γ)-Sgo1_1-155_ complex were pooled, concentrated, mixed with a 2-molar excess of CPC_I1-58SB_ and incubated for 30 min prior to injection. Individual PP2A(B56γ) and Sgo1_1-155_ were run as controls.

Interaction trials of the PP2A(B56γ)-CPC_I1-58SB_ complex, as well as those involving CPC_I1-58SBΔ110-188,_ Borealin_D120A/I123A/E125A_, Borealin_V116A/V119A/E121A_ and CPC_I1-58SB1-221_ were performed in a Superdex 200 Increase 10/300 column pre-equilibrated with 25 mM Hepes pH 8.0, 250 mM NaCl, 5 % glycerol and 2 mM DTT. 200 μg of PP2A(B56γ) were mixed with a two-molar excess of CPC_I1-58SB_, CPC_I1-58SBΔ110-188,_ Borealin_D120A/I123A/E125A_ or Borealin_V116A/V119A/E121A_, or with a six-molar excess of CPC_I1-58SB1-221_, and incubated for 30 min prior to injection. Individual proteins and the wild-type PP2A(B56γ)-CPC_I1-58SB_ complex (for the interaction trials with mutants and truncations) were run as controls.

For the interaction trials with isolated B56γ subunit, 300 μg of B56γ_WT_ or B56γ_R197E_ were mixed with equimolar amounts of CPC_I1-58SB_ and run in a Superdex 200 Increase 10/300 column pre-equilibrated with 25 mM Hepes pH 8.0, 150 mM NaCl, 5 % glycerol and 2 mM DTT. Individual proteins were run as controls.

For the interaction trials involving Sgo1_1-155 Nmut_, 35 μg of PP2A(B56γ) were mixed with a three-molar excess of Sgo1_1-155_ and 1.5-molar excess of CPCI1-58SB, incubated for 30 min and injected in a Superdex 200 Increase 5/150 column pre-equilibrated with 25 mM Hepes pH 8.0, 250 mM NaCl, 5 % glycerol and 2 mM DTT. The binary interactions and the wild-type ternary complex were used as controls.

### Cross-linking coupled to Mass-spectrometry

The PP2A(B56γ)-Sgo1_1-155_-CPC_I1-58SB_, PP2A(B56γ)-Sgo1_1-155_ and PP2A(B56γ)-CPC_I1-58SB_ complexes were subjected to SEC as described in ‘Interaction trials by SEC’. For the PP2A(B56γ)-Sgo1_1-155-_CPC_I1-58SB_ and PP2A(B56γ)-CPC_I1-58SB_ complexes, buffer containing 25 mM Hepes pH 8.0, 250 mM NaCl, and 2 mM DTT was used. For the PP2A(B56γ)-Sgo1_1-155_ complex, the cross-linking reaction buffer was phosphate saline buffer at pH 7.4.

Titrations of EDC and Sulfo-NHS (added at a 1:2.2 molar ratio, respectively), were performed to determine the optimal cross-linker concentration that yielded a cross-linked population of the expected molecular weight. Per cross-linking reaction, 10 μg of the protein complex were mixed with corresponding amounts of EDC and Sulfo-NHS and allowed to react for 90 min at room temperature in the dark. The reaction was quenched by adding Tris-HCl pH 8.0 at a final concentration of 50 mM. Crosslinked products were separated by SDS-PAGE using 4-12% Bis-Tris gels and NuPAGE MOPS SDS running buffer (Invitrogen), followed by staining with InstantBlue Coomassie protein stain (Expedeon). Samples crosslinked with 50 μg EDC (for the PP2A(B56γ)-Sgo1_1-155_-CPC_I1-58SB_ complex), 30 μg of EDC (for the PP2A(B56γ)-Sgo1_1-155_ complex), and 60 μg EDC (for the PP2A(B56γ)-CPC_I1-58SB_ complex) were processed for mass-spectrometry analysis.

Bands of interest were excised using a clean scalpel, cut into approximately 1 mm³ pieces and destained by incubating them in a 1:1 (v/v) mixture of 50 mM ammonium bicarbonate (ABC) and acetonitrile (ACN) for 1 h at 37 °C. Reduction was performed by incubating the gel pieces with 10 mM DTT in 50 mM ABC for 30 min at 37 °C, followed by alkylation with 55 mM iodoacetamide for 20 min at room temperature in the dark. Proteins were digested overnight at 37 °C using 13 ng/μl trypsin in 10 mM ABC containing 10% (v/v) ACN. The resulting peptide solution was acidified to pH 1-2 using 10% trifluoroacetic acid and loaded onto C18 StageTips (*49*).

Peptides were subsequently analyzed by liquid chromatography-tandem mass spectrometry (LC– MS/MS) using an Orbitrap Fusion Lumos mass spectrometer (Thermo Fisher Scientific) and data were analyzed as reported in Gireesh et al., 2025, Abad et al., 2022 and Abad et al., 2019 (*33–35*)

### Cryo-EM sample preparation

To prepare grids containing the PP2A(B56γ)-Sgo1_1-155_-CPC_I1-58SB_ complex, 100 μg of PP2A(B56γ) were mixed with 35 μg of Sgo1_1-155_ and 100 μg CPC_I1-58SB_ and injected onto a Superose 6 Increase 5/150 (Cytiva) equilibrated with 25 mM Hepes pH 8.0, 150 mM NaCl and 2 mM DTT. Fractions containing the stoichiometric complex were pooled, concentrated up to 0.9 mg/ml and crosslinked with 0.01 % glutaraldehyde for 5 min at 4 °C. The reaction was quenched with the addition of 50 mM Tris-HCl pH 8.0.

For the PP2A(B56γ)-Sgo1_1-155_ complex, 60 μg of PP2A(B56γ) were mixed with 20 μg of Sgo1_1-155_ and dialyzed overnight at 4 °C in buffer containing 25 mM Hepes pH 8.0, 100 mM NaCl and 2 mM DTT. Sample at a concentration of 0.8 mg/ml was crosslinked with 0.005 % glutaraldehyde for 5 min at 4 °C, reaction quenched with 50 mM Tris-HCl pH 8.0.

For the PP2A(B56γ)-CPC_I1-58SB_ complex, 100 μg of PP2A(B56γ) were mixed with 100 μg CPC_I1-58SB_ and dialyzed overnight against a buffer containing 25 mM Hepes pH 8.0, 200 mM NaCl and 2 mM DTT. Sample at a concentration of 1.2 mg/ml was crosslinked with 0.01 % glutaraldehyde for 5 min at 4 °C and the reaction was quenched with 50 mM Tris-HCl pH 8.0. 0.005 % Tween-20 was added prior to vitrification.

Grid preparation was performed using a Leica EM GP plunger working at 10 °C and 95% humidity. At all cases, samples were applied onto Quantifoil R2/2 300 mesh grids previously glow-discharged for 7 s in the presence of air using a current of 20 mA. Per grid, 4.5 μl of sample was applied and blotted for 2.5 s.

### Cryo-EM data processing and model building

Collection parameters are summarized in Table S1. All datasets were collected in a Titan Krios transmission electron microscope operating at 300 keV equipped with a Falcon 4i direct electron detector and a Selectris X Energy Filter (energy slit width 5 eV) (ThermoFisher). Microscope was working at a nominal magnification of x165,000 (resulting in a pixel size 0.727 Å/px), with a total dose of 40 e/Å^2^ per movie and variable exposure time to keep dose constant. Defocus range varied between -2.6 μm and -0.5 μm in 0.3 μm steps. Smart EPU software (ThermoFisher) was used for grid screening and automated data acquisition.

For the PP2A(B56γ)-Sgo1_1-155_-CPC_I1-58SB_ complex, a total of 16,787 movies were collected. The dataset was entirely processed using CryoSPARC v4.7 and v5.0 (*50*). Movies were subjected to patch motion correction and patch CTF estimation and only micrographs with an estimated resolution of 4 Å were used for downstream processing. Blob picker was initially used for particle picking in the entire dataset, yielding 20,838,210 particles extracted with a box size of 356 pixels with 3.7 times binning (2.6 Å/pixel). Following 2D classification, 106,349 particles showcasing secondary structure features were used to train Topaz (*51*). Iterative rounds of Topaz-based picking and 2D classification yielded 388,545 good particles that were re-extracted with an unbinned box size of 450 pixels (0.727 Å/pixel). Two rounds of ab initio reconstruction followed by heterogeneous refinement and one round of 2D classification were used to isolate a homogeneous particle population containing 216,558 particles. These particles were used as inputs for ab initio reconstruction followed by heterogeneous refinement with two classes, which yielded a class containing 122,387 particles with clear features of the PP2A(B56γ)-Sgo1_DH_ core with an occupied LxxIxE-binding pocket and additional Sgo1 and CPC-derived densities. Homogeneous refinement of this class yielded the ‘consensus’ map, with an estimated resolution of 2.89 Å. Non-uniform (NU) refinement (*52*) of these particles with a tight mask around the well-defined core yielded the ‘high-resolution’ map with an estimated resolution of 2.82 Å. The ‘consensus’ map particles were used as inputs of a 3D classification job with a mask around the PP2A A-B56 interface and four classes. NU-refinement of the class with a clear definition of the Borealin loop (31,663 particles) yielded the ‘Borealin loop’ map, with an estimated resolution of 3.14 Å.

For PP2A(B56γ)-Sgo1_1-155_, 10,182 movies were collected and subjected to motion correction using MotionCor2 (*53*). Micrographs were subsequently processed in CryoSPARC v4.7 and v5.0. Patch CTF estimation was performed, and initial particle picking was done using blob picking on 256 micrographs. Particles were extracted with a box size of 356 px and 3.7 times binning (2.6 Å/pixel). Following 2D classification, 10,338 particles were used as templates for template-based picking in a subset of 640 micrographs. After several rounds of 2D classification, 21,076 particles with clear secondary structure features were selected and used to train Topaz. Iterative rounds of Topaz-based picking combined with 2D classification yielded 273,925 good particles that were subsequently re-extracted with an unbinned box size of 400 pixels (0.727 Å/pixel). Two rounds of ab initio reconstruction combined with heterogeneous refinement with two classes each were performed to isolate 79,230 particles that yielded a clear reconstruction of the PP2A(B56γ)-Sgo1_DH_ core with additional Sgo1-derived densities. Homogeneous refinement of these particles yielded a map at a resolution of 3.26 Å resolution that was used as input for masked 3D classification and NU-refinement. NU-refinement of these particles with a tight mask around the core yielded the ‘high-resolution’ map, with an average resolution of 3.11 Å. 3D classification with 3 classes and a mask above Sgo1_DH_ followed by NU-refinement of the class showcasing clear Sgo1 N-terminal helices (27,289 particles) resulted in the ‘consensus’ map, with an average resolution of 3.78 Å. 3D classification with 3 classes and a mask around Sgo1 C-terminal region yielded 3 classes with peptide-like densities corresponding to Sgo1_IDR_.

For the PP2A(B56γ)-CPC_I1-58SB_ complex, 10,356 movies were collected. MotionCor2 was used for motion-correction, with further processing done in CryoSPARC v4.7 and v5.0. Micrographs were subjected to patch CTF estimation, and exposure curation was performed with a CTF resolution estimation cutoff of 4 Å. Initial particle picking was performed with blob picking over the entire dataset (extraction box size of 356 px with a binning of 3.7 - 2.6 Å/pixel). Several rounds of 2D classification resulted in 48,675 particles that were used to train Topaz. After 2D classification, particles were separated into common views (42,221 particles) and rare views (12,278 particles), with Topaz models generated for both and used for subsequent picking. Resulting particles were merged, duplicates were removed, and several rounds of 2D classification were performed. 153,131 particles showcasing clear secondary structure features were used for ab initio reconstruction followed by heterogeneous refinement with two classes each. 112,953 particles from the best class were re-extracted at an unbinned box size of 356 pixels (0.727 Å/pixel), and further 2D classification, NU-refinement and local CTF refinement was performed. Ab initio reconstruction followed by heterogeneous refinement with two classes each yielded a class with clear PP2A(B56) features and the LxxIxE-binding pocket occupied (83,114 particles). 2D classification followed by NU-refinement resulted in the ‘high-resolution’ map, with an estimated resolution of 3.05 Å (71,070 particles). 3D classification with 4 classes and a mask around the A-B56 interface and the B56 C-terminal region resulted in a class, containing 16,941 particles, with clear features for the Borealin loop and dimerization domain. Further NU-refinement yielded the ‘Borealin loop and DD’ map at 3.51 Å overall resolution.

Initial rigid body fitting with AlphaFold3-predicted (*54*) PP2A(B56γ)-Sgo1_51-106_-Borealin_120-127_, PP2A(B56γ)-Sgo1_51-106_ and PP2A(B56γ)-Borealin_120-127_ into the ‘high-resolution’ maps was done in ChimeraX 1.12 (*55*). To obtain the final models, iterative rounds of real-space refinement using Phenix (v1.21.2) (*56*) and Coot-1 (*57*) were performed. For the PP2A(B56γ)-CPC_I1-58SB_ complex, rigid body fitting of the Borealin_DD_ NMR structure (PDB: 2KDD) was done in ChimeraX 1.12. Real space refinement was done with Namdinator (*58*) against the Borealin loop and DD map.

### Multiple-sequence alignment

Multiple sequence alignment of Borealin from Homo sapiens (Hs), Sus scrofa (Ss), Bos taurus (Bt), Equus caballus (Ec), Balaenoptera musculus (Bm), Rattus norvegicus (Rn), Mus musculus (Mm), Gallus gallus (Gg), Danio rerio (Dr) and X. laevis (Xl) was performed using Clustal Omega (*59*) with default settings. Analysis and visualization was done in JalView 2.11.5.3 (*60*)

### Computational modelling and Molecular Dynamics simulations

The PP2A(B56γ)-Sgo1 model was generated using AF3x (*43*), an AlphaFold3-based method that incorporates XL-MS-derived restraints through explicit linker modelling. Predictions were generated using two model seeds, with one diffusion sample per seed. The highest-ranked model was selected according to the AlphaFold3 ranking score.

The selected model was docked into the consensus PP2A(B56γ)-Sgo1 cryo-EM map (Fig. S3). Two possible arrangements of the N-terminal helices were considered, corresponding to cis (within-protomer) and trans (between-protomer) configurations of the K19-E77 crosslink. Predicted helices (residues 11-38) were fitted as rigid bodies to the density by maximizing the mean map value at backbone atoms. The trans arrangement was used for subsequent molecular dynamics simulations. Sgo1 residues 2-10 and 39-47 were rebuilt using MODELLER 10.8 (*61*). The resulting model was refined using OpenMM 8.6.1 with the Amber ff14SB force field and GBn2 implicit solvent (*62–64*). Only the rebuilt atoms were allowed to move. A density potential, −kΣρ (k = 2.0 kJ mol⁻¹ σ⁻¹), was applied to regions of the cryo-EM map not accounted for by the fixed model. Refinement consisted of energy minimization, heating to 300 K, 120 ps of simulated annealing (300-360-300 K), followed by a final energy minimization. Crosslink satisfaction was assessed using a Cα-Cα distance threshold of 20 Å. This criterion was used to assess agreement with the XL-MS data.

The refined trans PP2A(B56γ)-Sgo1 model was converted to the Martini 3 force field (v3.0.0) using martinize2 (vermouth 0.15.0) (*65, 66*). Secondary structure was assigned from the atomistic model using DSSP 4.6.1 (*67, 68*). An intrachain elastic network was generated between backbone (BB) beads separated by 0.5-0.9 nm, with a force constant of 700 kJ mol⁻¹ nm⁻² (*69*). Inter-subunit interfaces were not constrained by elastic bonds and were instead maintained through non-bonded interactions. Within each Sgo1 protomer, separate elastic network units were defined to allow relative movement of the N-terminal helices and coiled coil. The protein was placed in a rhombic dodecahedral box with a box vector of 19.45 nm and solvated with Martini water and 0.15 M NaCl, including neutralizing ions. Energy minimization was performed using the steepest-descent algorithm until the maximum force was below 100 kJ mol⁻¹ nm⁻¹.

All molecular dynamics simulations were performed using GROMACS 2026.3 with the Verlet cutoff scheme (*70*). Electrostatic interactions were treated using the reaction-field method (relative dielectric constant, εr = 15) (*71, 72*), and Lennard-Jones interactions were calculated using a 1.1 nm cutoff with a potential-shift modifier. Protein and solvent were coupled separately to a V-rescale thermostat at 310 K (τT = 1 ps) (*73*). Bond constraints were enforced using LINCS.

For the PP2A(B56γ)-Sgo1 complex, each replica underwent an initial 1 ns equilibration with a 5 fs time step and backbone position restraints (1,000 kJ mol⁻¹ nm⁻²), followed by 5 ns of equilibration with a 20 fs time step and no position restraints. Equilibration was performed at 1 bar using the C-rescale barostat (*74*). Production simulations were performed using a 20 fs time step and the Parrinello-Rahman barostat (*75*). Five independent replicas were initiated from the same energy-minimized structure. No crosslink bonds or crosslink-derived distance restraints were included during energy minimization, equilibration or production simulations.

For the PP2A(B56γ)-Sgo1 complex, frames were sampled every 100 ps and aligned to the starting structure using the Kabsch algorithm in the cryo-EM reference frame (*76*). Internal RMSD was calculated separately for the PP2A(B56γ) core and Sgo1 dimerization helix (Sgo1_DH_). Sgo1_DH_ displacement was measured following alignment on PP2A(B56γ), while retention of the N-terminal arrangement was assessed by calculating the RMSD of Sgo1 residues 2-47 after alignment on the Sgo1_DH_. BB-BB distances were monitored for cross-link pairs. Contacts were defined by a bead-pair distance below 6 Å. For each replica, mean values were calculated over the final 100 ns of the trajectory.

Coarse-grained trajectories of the PP2A(B56γ)-Sgo1 complex were back-mapped at 1 ns intervals by local template fitting. Frames were sampled every 10 ns from the first 1 µs of replicas 1-5 and energy-minimized in OpenMM 8.6.1 using Amber ff14SB and GBn2 implicit solvent(*62–64*). Occupancy maps for Sgo1 residues 2-47 and 107-155 were calculated from unminimized back-mapped trajectories over the first 1 µs of replicas 1-5 (1,001 frames per replica). Maps were generated on a 1 Å grid in the cryo-EM reference frame. A grid point was considered occupied if any heavy atom of the corresponding protein segment was located within 2.0 Å. Occupancy was calculated as the fraction of frames in which each grid point was occupied (0–1), both independently for each replica and across pooled replicas. The resulting maps were compared with the experimental cryo-EM density at a threshold of 3σ.

The ternary PP2A(B56γ)-Sgo1-CPC complex was modelled by combining published structural information for the PP2A(B56γ)-Sgo1 complex (PDB: 3FGA) and the CPC triple-helical bundle (PDB: 2QFA) with the consensus cryo-EM density map of the PP2A(B56γ)-Sgo1-CPC complex. Structural models were assembled and fitted into the cryo-EM map using Coot (*57*), followed by real-space refinement in PHENIX (*56*). The refined ternary complex model was converted to the Martini 2.2 force field using CHARMM-GUI. Secondary structure was assigned from the atomistic model using DSSP (*67, 68*). An elastic network was generated using ElNeDyn to preserve the assigned secondary structure (*69*). Inter-domain elastic connections within individual proteins were removed to avoid artificial constraints between domains. The complex was placed in a cubic box with a side length of 19.98 nm and solvated using water and 0.15 M NaCl. Two rounds of steepest-descent energy minimization were performed until the maximum force was below 100 kJ mol⁻¹ nm⁻¹.

The energy-minimized system was equilibrated for 100 ns using a 20 fs time step and backbone position restraints. Production simulations were subsequently performed for 1 µs without position restraints, using the V-rescale thermostat and Parrinello-Rahman barostat (*73, 75*).

The volume sampled by the ternary PP2A(B56γ)-Sgo1-CPC complex during molecular dynamics simulations was calculated using a custom Python script based on MDAnalysis, NumPy and SciPy (*77–79*). Trajectory frames were sampled every 10 ns, and a Gaussian filter with a 3 Å standard deviation was applied to the resulting volume to generate maps comparable to the experimental cryo-EM density. Volumes were calculated for the complete complex, aligned on the PP2A A subunit, and separately for each CPC. The resulting volumes were fitted to the experimental cryo-EM map using ChimeraX, and cross-correlation values were calculated. Trajectory analyses were performed using MDAnalysis 2.10, MDTraj 1.11, NumPy and mrcfile 1.5.4 (*77, 78, 80, 81*), together with custom Python scripts ([GitHub repository to be provided]). Structural visualization was performed using PyMOL 3.1 (Schrödinger, LLC) and ChimeraX (*55*).

### Rescue experiments and immunofluorescence microscopy with Hela Kyoto cells

The pCDNA3.1 vector containing N-terminally tagged 3xMyc-Borealin was a gift from E. Nigg’s laboratory (Biozentrum, The Centre for Molecular Life Sciences, Basel, Switzerland). Mutations of Borealin were obtained using the Quikchange site-directed mutagenesis method (Stratagene). The oligonucleotides targeting the 3′ UTR of Borealin (5′-AGGTAGAGCTGTCTGTTCAdTdT-3′) or targeting luciferase as a control (5′-CGUACGCGGAAUACUUCGAdTdT-3′) were described previously (*82*). All siRNA oligonucleotides were purchased from Qiagen. Cells were transfected using jetPRIME (Polyplus Transfection) as previously described (*83*). 50 nM siRNA oligonucleotides and 300 ng of Myc-Borealin vectors were used for transfections. Cells were fixed in 4% PFA 36 h after transfection.

The following antibodies were used for indirect immunofluorescence: anti-myc (1:200; 9E10; Merck Millipore), anti-Survivin (1:500; NB500-201; Novus), and anti-ACA (1:300; 15-235; Antibodies Inc.). The secondary antibodies used were FITC-conjugated AffiniPure donkey anti-mouse IgG (1:300; Jackson Immunoresearch), TRITC-conjugated AffiniPure goat anti-rabbit (1:300; Jackson Immunoresearch), Cy5-conjugated AffiniPure donkey anti-human (1:400; Jackson Immunoresearch). DAPI was used for DNA staining. Imaging was performed at room temperature using a Nikon Ti2 microscope with Photometrics Prime 95B camera and 100× NA 1.4 CFI Plan Apo Lambda D objective with oil immersion (refractive index = 1.514) using the NIS Elements (Nikon) software. The 0.2 µm spaced z-stacks were deconvolved using Huygens (Scientific Volume Imaging) software. Shown images are maximum-intensity projections.

For quantification of the centromeric levels of Survivin, the acquired images were deconvolved using Huygens (Scientific Volume Imaging) software, and the centromere intensity of Survivin was quantified using an ImageJ plugin (Zenodo). Briefly, the plugin quantifies the maximum fluorescence signal of Survivin in a 10×10 pixel box immediately outside the centromere, defined with the ACA staining and values obtained after subtracting the background from the maximum intensities in the square area were averaged and normalized to the mean of ACA intensity. To compare data from different replicates, values were averaged and normalized to the mean of Survivin/ACA intensity in the Myc-Borealin rescue condition. Statistical significance of the difference between normalized intensities at the centromere region was established by a Mann–Whitney *U* test using Prism 11.

Quantification of anaphases displaying chromosome bridges or lagging chromosomes was performed 48 h after HeLa Kyoto cells were transfected. The experiment was performed in 5 repeats, and a minimum of 100 anaphases were quantified per replicate. For the Monastrol assay, HeLa Kyoto cells were synchronized with 100 μM Monastrol for 16 h and released into 5 μM MG132 for 1 hour. Observed metaphases were classified as aligned, and misaligned. The experiment was performed in triplicate, and a minimum of 155 metaphases were quantified per replicate.

### Immunoprecipitation

Doxycycline-inducible HeLa cell line expressing YFP-PP2A-B56γ (kindly provided by the A. Saurin laboratory) were used for immunoprecipitation experiments. Cells were transfected with either Myc-Borealin wild-type (WT) or Myc-Borealin 3A expression constructs using JetPRIME (Polyplus) according to the manufacturer’s instructions. The following day cells were treated with 1 µM palbociclib for 18 h, released for 8h and subsequently treated with 50 ng/mL nocodazole and 1 µg/mL doxycycline. Cells were incubated at 37°C for 16 h to induce Borealin expression and achieve mitotic arrest. To enrich in prometaphase/metaphase, cells were then released from nocodazole arrest for 45 min before harvesting. Cells were then lysed in lysis buffer (50 mM Tris-HCl (pH 7.4), 150 mM NaCl, 40 mM β-glycerophosphate, 0.5% (v/v) IGEPAL CA-630, 0.1% sodium deoxycholate, protease and phosphatase inhibitor cocktail, 100 µM ATP, 100 µM MgCl₂ and 0.5 mM EDTA). Cell lysates were incubated for 30 min at 4°C with gentle rocking and subsequently sonicated using a Bioruptor Pico sonicator (30 s on/30 s off, 12 cycles, at 4°C). Lysates were centrifuged at 13,000 rpm for 10 min at 4°C to remove insoluble material. Lysates were incubated with GFP-Trap magnetic agarose beads (ChromoTek) for 1 h at 4°C. After a lysis buffer wash, proteins were eluted with 2× Laemmli SDS sample buffer and subsequently analysed by SDS-PAGE followed by western blot. The antibodies used for the western blot were mouse anti-myc (clone 9E10; Merck), rabbit anti-GFP (Abcam) and mouse anti-PP2A-C (clone 1D6; Millipore). Secondary antibodies used were IRDye 800CW donkey anti-mouse IgG, IRDye 800CW donkey anti-rabbit IgG, IRDye 680RD goat anti-mouse IgG (LI-COR Biosciences).

### hTERT-RPE1 cell culture, cell-line generation and immunofluorescence microscopy

hTERT-RPE1 cells were cultured in DMEM supplemented with 10% fetal bovine serum (FBS) and 10 mM HEPES (pH 7.5) at 37 °C in a humidified atmosphere containing 5% CO₂.

The OsTIR1(F74G)-expressing Borealin-mAID-mClover RPE1 cell line and the doxycycline-inducible Borealin-mCherry WT replacement cell line were generated as previously described in Gireesh et al., 2025 (*35*). To generate the doxycycline-inducible Borealin-mCherry 3A replacement cell line, the Borealin 3A mutant was introduced into the hROSA26 locus of Borealin-mAID-mClover cells using CRISPR–Cas9-mediated homologous recombination, following the same procedure as described previously (Gireesh et al., 2025). The Borealin 3A mutant contained the D120A, I123A, and E125A substitutions.

For Borealin replacement, expression of Borealin-mCherry WT or 3A was induced by treatment with 1 μg/mL doxycycline for 12 h, followed by depletion of endogenous Borealin-mAID-mClover with 1 μM 5-Ph-IAA for 1 h.

hTERT-RPE1 cells grown on coverslips were fixed with 2% paraformaldehyde in PBS for 10 min and permeabilized with 0.2% Triton X-100 in PBS for 10 min. After blocking with 3% BSA in PBS for 10 min, cells were incubated with primary antibodies diluted in 3% BSA in PBS. The following primary antibodies were used: anti-CENP-A (1:500; D115-3, MBL) and anti-mCherry (1:1000; GTX128508, GeneTex). After incubation with secondary antibodies and counterstaining of DNA with 0.1 μg/mL DAPI, cells were mounted using ProLong Gold Antifade Mountant (Invitrogen).

Images were acquired using a Plan-Apochromat 63×/1.4 NA oil-immersion objective on an LSM 880 confocal laser-scanning microscope (Zeiss). Borealin-mCherry was detected by immunofluorescence using an anti-mCherry antibody. ROI-based measurements were obtained from maximum-intensity projection images. ROIs with a diameter of 14 pixels were centered on Borealin-mCherry signals, whereas 10-pixel-diameter ROIs were centered on CENP-A signals. Image processing and intensity measurements were performed using Fiji (ImageJ). For each cell, background-subtracted anti-mCherry intensities were normalized to the mean background-subtracted CENP-A intensity measured in the same cell.

For the analysis of chromosome congression and segregation defects. hTERT-RPE1 cells were treated with 100 μM monastrol for 3 h and released from the arrest. At 40 min after release, cells were fixed and stained with DAPI. At this time point, chromosome congression defects were assessed in metaphase cells based on the presence of one or more unaligned chromosomes, whereas chromosome segregation defects were assessed in anaphase cells based on the presence of lagging chromosomes and chromosome bridges.

### Western blot

To study the expression levels of each of the Borealin constructs, HeLa Kyoto cells were transfected in 6-well dishes using 100 nM siRNA oligonucleotides and 600 ng of Myc-Borealin vectors. Cells were treated with Palbociclib for 18 hours, released for 8 hours and then incubated with nocodazole for 12 hours. After nocodozole treatment, cells were released for an hour before harvesting. Cells were lysed in lysis buffer (50 mM Tris, pH 7.4, 400 mM NaCl, 40 mM β-glycerol phosphate, 10 mM NaF, 0.5% IGEPAL, 0.1% deoxycholate, 100 µM ATP, 100 µM MgCl_2_, and protease inhibitor cocktail tablet (Roche Diagnostics); adapted from (*83*). Lysates were then sonicated and spun down at 15,000 *g* for 15 min and Laemmli buffer was added to the supernatant. This sample was boiled for 5 min, and analyzed by SDS-PAGE followed by Western blotting. The antibodies used for the immunoblot were mouse anti-tubulin (1:10,000; T5168; Sigma Aldrich), and mouse anti-Borealin (1:1,000; M147-3; MB). Secondary antibodies used were goat anti-mouse 680 (LI-COR) at 1:5,000 dilution. Immunoblots were imaged using the Odyssey CLx system. The following primary antibodies were used for western blotting of hTERT-RPE1 cell lines: Anti-Borealin (1:1000; M147-3, MBL); Anti-mCherry (1:1000; GTX128508, GeneTex); Anti-α-Tubulin (1:5000; clone B-5-1-2, #T6074, Millipore).

### Live cell imaging

For time-lapse microscopy, HeLa cells were plated in 35mm glass bottom dishes (matTek corporation) and transfected using 100 nM siRNA oligonucleotides and 600 ng of Myc-Borealin vectors. After 48 hours, DNA was stained using SPY650-DNA Live cell DNA stain (Spirochrome) an hour before imaging. Cells were imaged using a Nikon Ti2 CSU-W1 Spinning Disk Live Imaging Microscope equipped with a 20 × /0.75 air CFI Plan Apochromat Lambda objective. During imaging, the atmosphere was maintained at a temperature of 37 °C and 5% CO_2_. Images were captured at 5-min intervals for 22 h at multiple positions with 50 ms exposure times at each time point. NIS Elements software (Nikon) was used to collect the data and analysis was done using ImageJ software.

To analyze mitotic progression in hTERT-RPE1, live-cell imaging was performed using a CQ1 confocal quantitative image cytometer (Yokogawa). Cells were cultured overnight in 24-well glass-bottom plates (Iwaki). DNA was stained with 100 nM SiR-DNA (Cytoskeleton, Inc.) for 1 h before image acquisition. Images were acquired every 1 min using a ×40 objective lens. Maximum-intensity projections of five Z-sections acquired at 1.5-μm intervals were used for analysis. For data analysis, images were processed using Fiji (ImageJ). Mitotic duration was measured from nuclear envelope breakdown (NEBD) to anaphase onset.

### Statistical analysis

Statistical analyses were performed using GraphPad Prism 10 (GraphPad Software). For analysis of chromosome segregation errors, chromosome misalignment and mitotic duration from NEBD to anaphase onset, statistical significance was assessed using either Mann-Whitney U test or a paired two-tailed t-test. For quantification of centromeric Borealin levels, the mean normalized Survivin or Borealin-mCherry intensity for each cell was used as the statistical unit, and WT and 3A cells were compared using either a Mann-Whitney U-test or an unpaired two-tailed Student’s t-test.

## Acknowledgments

We thank the Wellcome Discovery Research Platform for Hidden Cell Biology, and we gratefully acknowledge support from the Microscopy, Proteomics, and Bioinformatics cores. We thank Adrian Saurin for sharing HeLa cell lines expressing YFP-PP2A-B56γ subunit (*19*). We also thank Karl-Peter Hopfner and his lab members for sharing resources and useful discussions.

## Funding

The research of A.A.J and his team was supported by the Wellcome Trust (Discovery Award 309153/Z/24/Z) and the European Union (ERC, CHROMSEG, grant no. 101054950).

The views and opinions expressed are, however, those of the authors only and do not necessarily reflect those of the European Union or the European Research Council. Neither the European Union nor the granting authority can be held responsible for them. The Wellcome Centre for Cell Biology is supported by core funding from the Wellcome Trust (grant 203149) and instrument grant (091020). This work was supported by funding for the Wellcome Discovery Research Platform for Hidden Cell Biology (226791). R.-S.N. and T.H. are supported by the Japan Society for the Promotion of Science (JSPS) through the following Grants-in-Aid for Scientific Research: 24H02286 (R.-S.N.), 24H02283 and 22H04996 (T.H.). V.D. is funded by the Darwin Trust of Edinburgh. P.N. is funded by EastBio (The Eastern Scotland Biosciences Doctoral Training Partnership). This work utilised the Gene Center cryo-EM facility funded by LMU Munich and DFG (Deutsches Forschung Gemeinschaft), the Wellcome Centre for Cell Biology Centre Optical Instrumentation Laboratory (COIL) / Light Microscopy Core (LMC) and Proteomics and Bioinformatics Core Platforms.

## Author contributions

Conceptualization: A.A.J.; Funding acquisition: A.A.J., T.H.; Investigation: P.P.S., V.D., R.-S.N., M.A.A., P.N., Z.Y., Z.F., J.Z., C.S., L.F.A., A.A.J.; Methodology: P.P.S., V.D., R.-S.N., M.A.A., P.N., Z.Y., Z.F., J.Z., C.S., L.F.A., A.A.J.; Software: Z.Y., Z.F., Writing – original draft: P.P.S., M.A.A., A.A.J.; Writing – review & editing: P.P.S., M.A.A., A.A.J., R.-S.N., V.D., Z.F.

## Competing interests

The authors declare no competing interests.

## Data, code, and materials availability

Data for the XL-MS experiments were deposited at PRIDE. Cryo-EM maps and structures have been deposited in the Protein Data Bank (PDB) and Electron Microscopy Database (EMDB) with accession numbers 50OA / EMD-100801 (PP2A(B56)-Sgo1), 50NZ / EMD-100800 (PP2A(B56)-CPC) and 50OC / EMD-100802 (PP2A(B56)-Sgo1-CPC). The ImageJ plugin was deposited in Zenodo.

---

**Figure S1:**
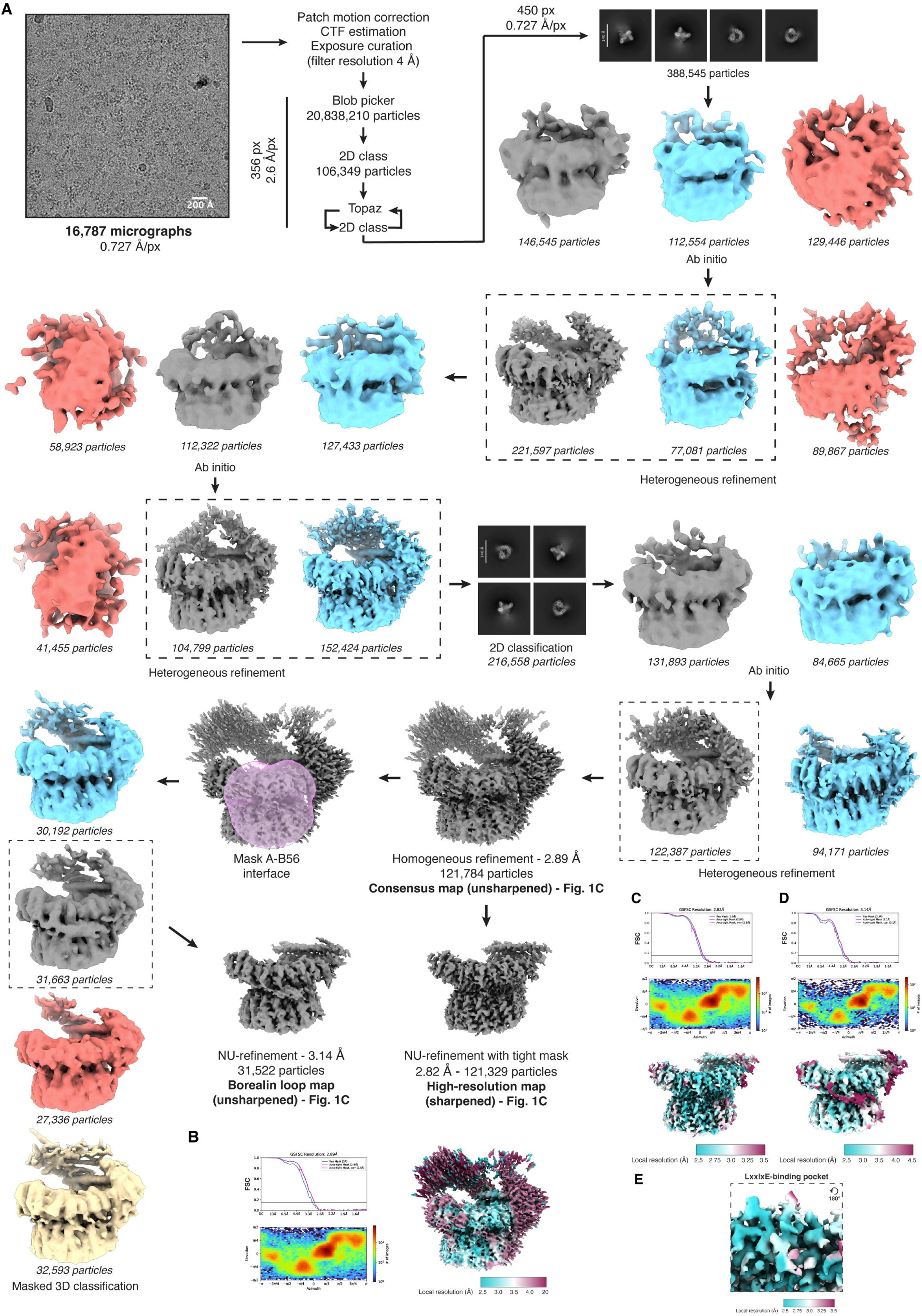
Cryo-EM processing workflow of the PP2A(B56)-Sgo1-CPC complex and map validation plots. **(A)** Cryo-EM processing workflow, **(B)** FSC curve, Azimuth plot and local resolution estimation of the ‘consensus’ map, **(C)** FSC curve, Azimuth plot and local resolution estimation of the ‘high-resolution’ map, (**D)** FSC curve, Azimuth plot and local resolution estimation of the ‘Borealin loop’ map. **(E)** Local resolution estimation of the LxxIxE-binding pocket of the ‘high-resolution’ map.

**Figure S2:**
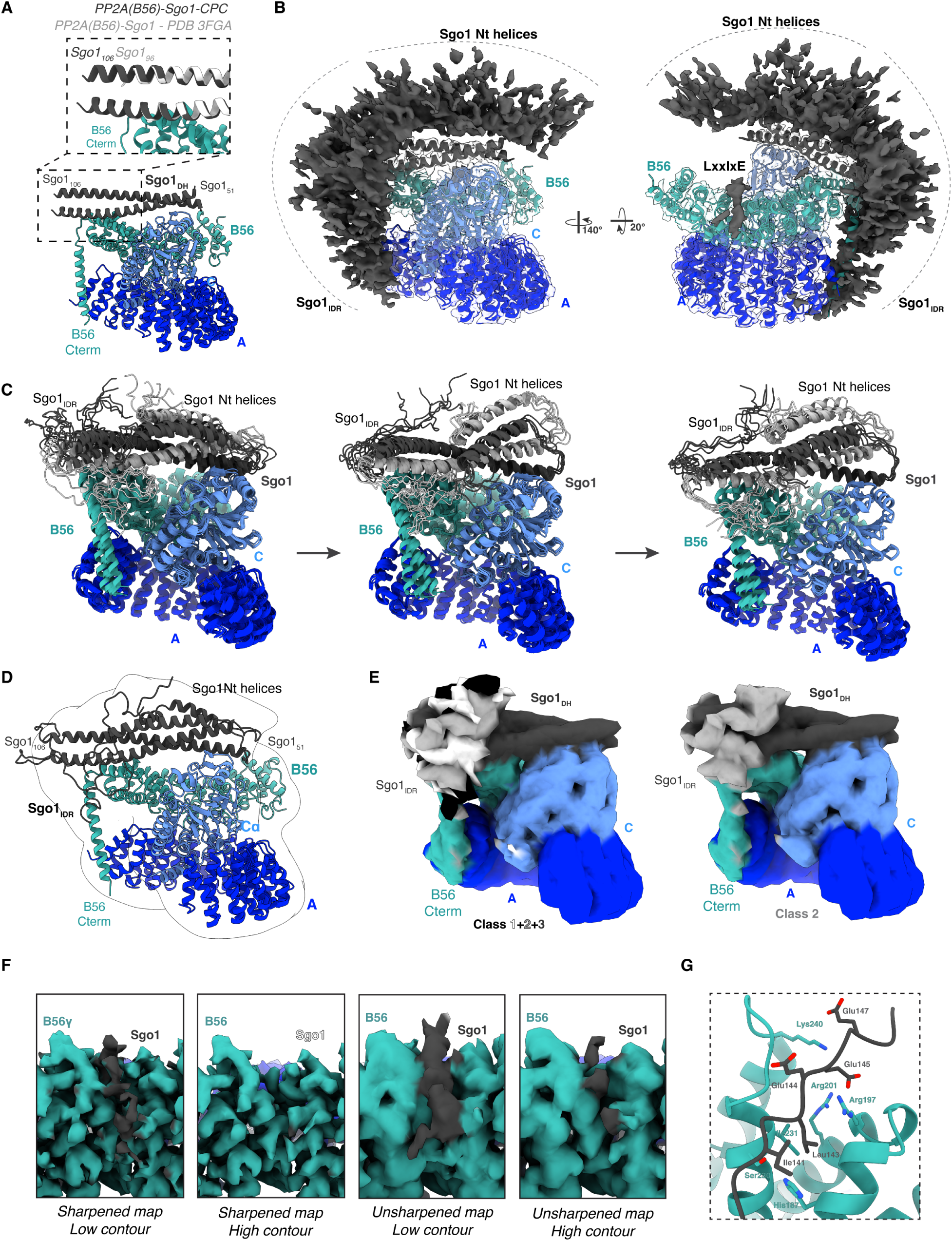
Structural characterization of the PP2A(B56)-Sgo1 complex. **(A)** Model for PP2A(B56)-Sgo1_DH_ in cartoon representation with a close-up view of the C-terminal end of the Sgo1_DH_ superposed against the Sgo1_DH_ from the PP2A(B56)-Sgo1 crystal structure (PDB 3FGA, shown in white). **(B)** Overlay of the ‘consensus’ and ‘high-resolution’ cryo-EM maps of the PP2A(B56)-Sgo1 complex (Fig. S3) in two orientations. Conformationally heterogeneous densities corresponding to Sgo1_IDR_ and Nt helices have been shown in dark grey and labelled accordingly. The PP2A(B56)-Sgo1_DH_ model is shown in cartoon representation, with corresponding density shown in transparency. **(C)** Selected frames of the MD trajectories for the PP2A(B56)-Sgo1 complex. Arrows represent the temporal progression of the trajectory. **(D)** Selected MD simulation frame of the PP2A(B56)-Sgo1 complex fitted into the gaussian-blurred ‘consensus’ map. **(E)** Overlay of the 3D classes of the PP2A(B56)-Sgo1 complex showcasing the Sgo1_IDR_ as peptide-like densities (left, Fig. S3). Sgo1_IDR_ for classes 1, 2 and 3 is shown in white, grey and black color, respectively. On the right, representative class 2 is shown. **(F)** Close-up views of the LxxIxE-binding pocket in the PP2A(B56)-Sgo1 complex. Sharpened and unsharpened maps with different contour levels are shown. **(G)** Close-up view of the AF3 model with Sgo1_141-147_ engaging the LxxIxE-binding pocket. Relevant residues have been shown in atomic representation.

**Figure S3:**
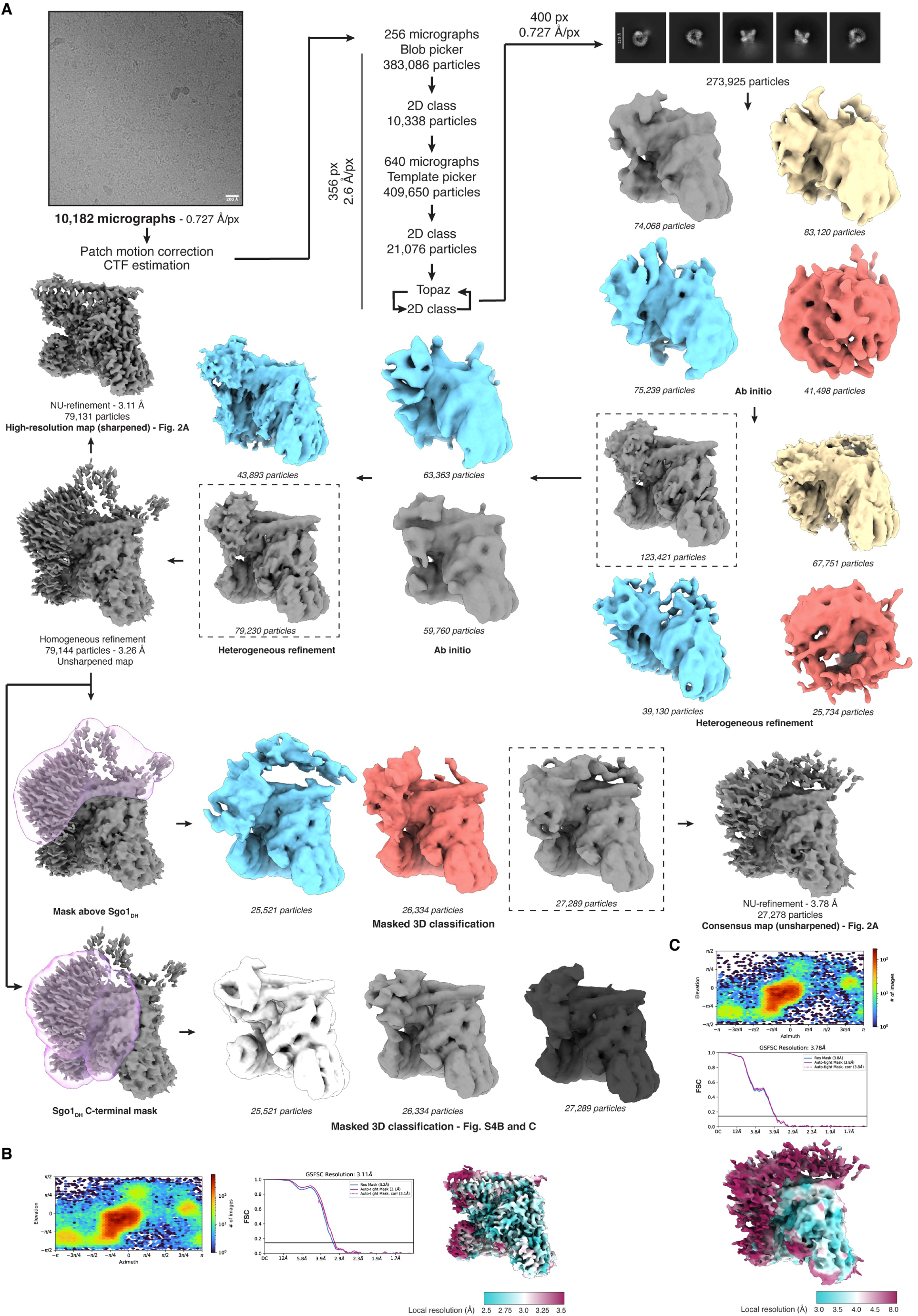
Cryo-EM processing workflow of the PP2A(B56)-Sgo1 complex and map validation plots. **(A)** Cryo-EM processing workflow, **(B)** Azimuth plot, FSC curve and local resolution estimation of the ‘high-resolution’ map, **(C)** Azimuth plot, FSC curve and local resolution estimation of the ‘consensus’ map.

**Figure S4:**
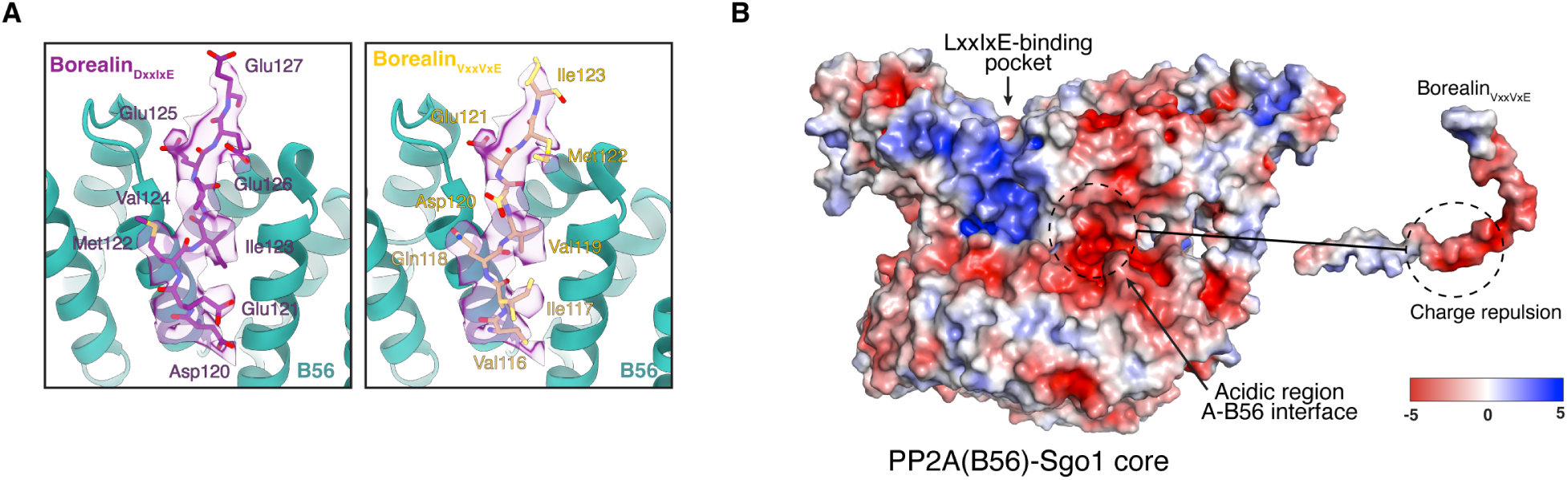
Interaction of Borealin loop containing LxxIxE-like motif and PP2A(B56). **(A)** Density fitting of the DxxIxE (right) and VxxVxE (left) peptides onto the PP2A(B56)-Sgo1-CPC_I1-58SB_ complex LxxIxE-binding pocket cryo-EM density (shown in purple with transparency). **(B)** APBS surface electrostatic potential of the PP2A(B56)-Sgo1 core (left) and the Borealin loop containing the VxxVxE motif (right). Negative charge is depicted as red, positive charge is depicted as blue and no charge is depicted in white. Regions that would generate charge repulsion have been indicated.

**Figure S5:**
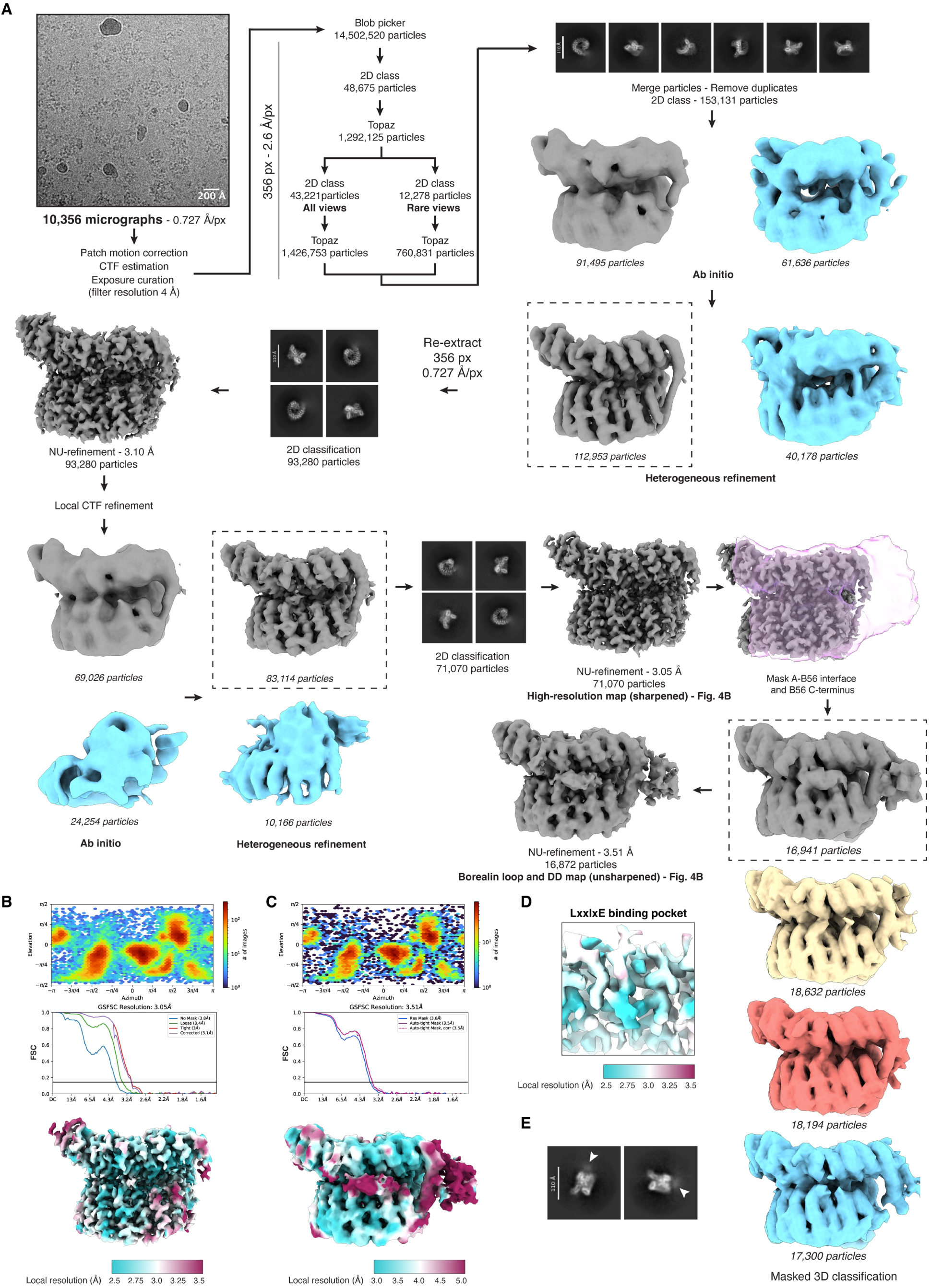
Cryo-EM processing workflow of the PP2A(B56)-CPC complex and map validation plots. **(A)** Cryo-EM processing workflow, **(B)** Azimuth plot, FSC curve and local resolution estimation of the ‘high-resolution’ map, **(C)** Azimuth plot, FSC curve and local resolution estimation of the ‘Borealin loop and dimerization domain’ map. **(D)** Local resolution estimation of the LxxIxE-binding pocket of the ‘high-resolution’ map. **(E)** Selected 2D classes showcasing the Borealin dimerization domain.

**Figure S6:**
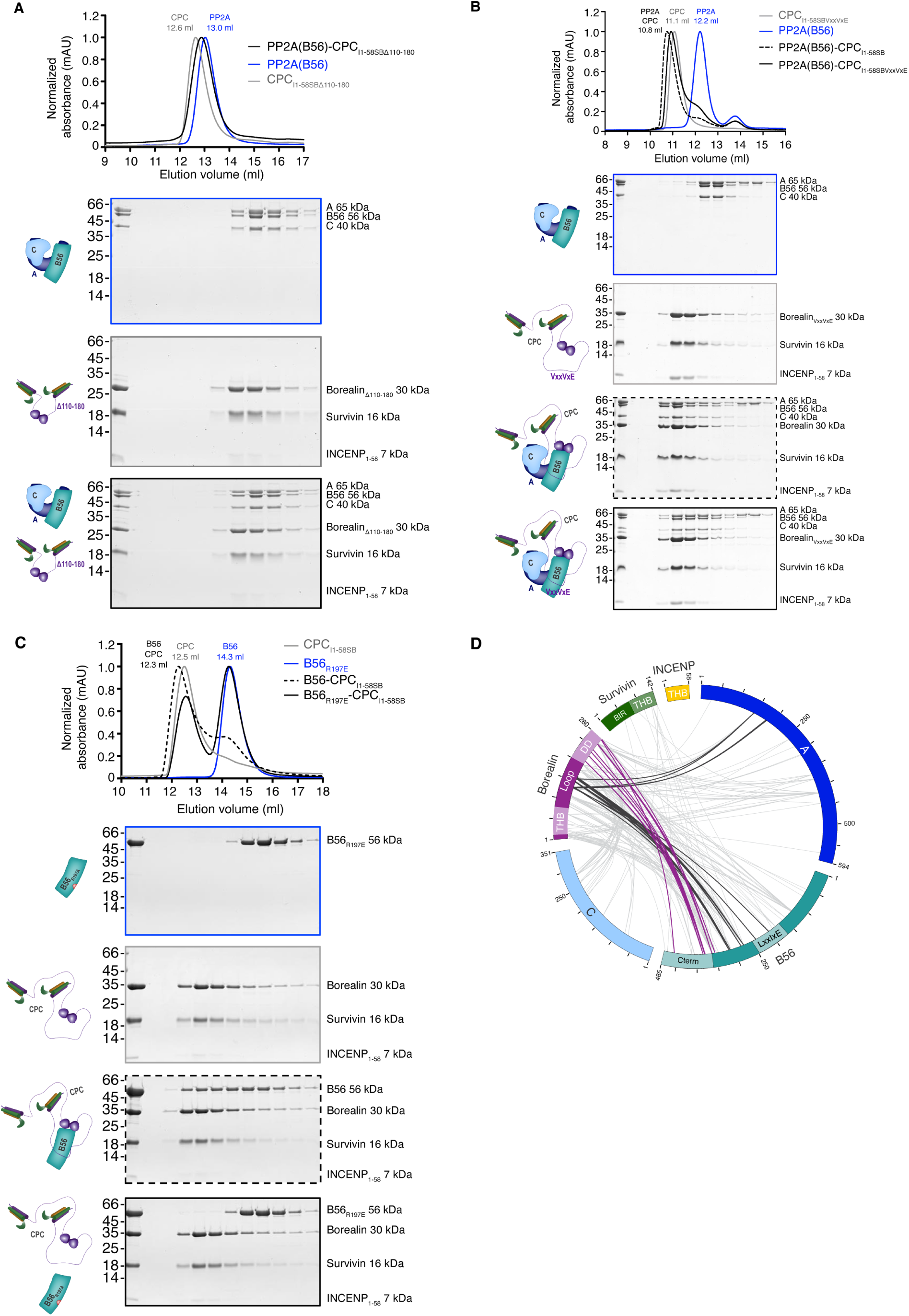
Biochemical validation of the requirement of Borealin loop for PP2A(B56) binding. **(A)** SEC profiles and corresponding SDS-PAGEs of PP2A(B56) (blue), CPC_I1-58SBΔ110-180_ (grey) and PP2A(B56)-CPC_I1-58SBΔ110-180_ (black). **(B)** SEC profiles and corresponding SDS-PAGEs of PP2A(B56) (blue), CPC_I1-58SBVxxVxE_ (grey), the PP2A(B56)-CPC_I1-58SB_ complex (dashed black) and PP2A(B56)-CPC_I1-58SBVxxVxE_ (solid black). **(C)** SEC profiles and corresponding SDS-PAGE of B56 subunit (blue), CPC_I1-58SB_ (grey), B56-CPC_I1-58SB_ (dashed black) and B56_R197E_-CPC_I1-58SB_ (solid black). **(D)** XL-MS circular diagram of the PP2A(B56)-CPC_I1-58SB_ complex, with cross-links involving the Borealin loop and Dimerization domain colored in dark grey and purple, respectively.

**Figure S7:**
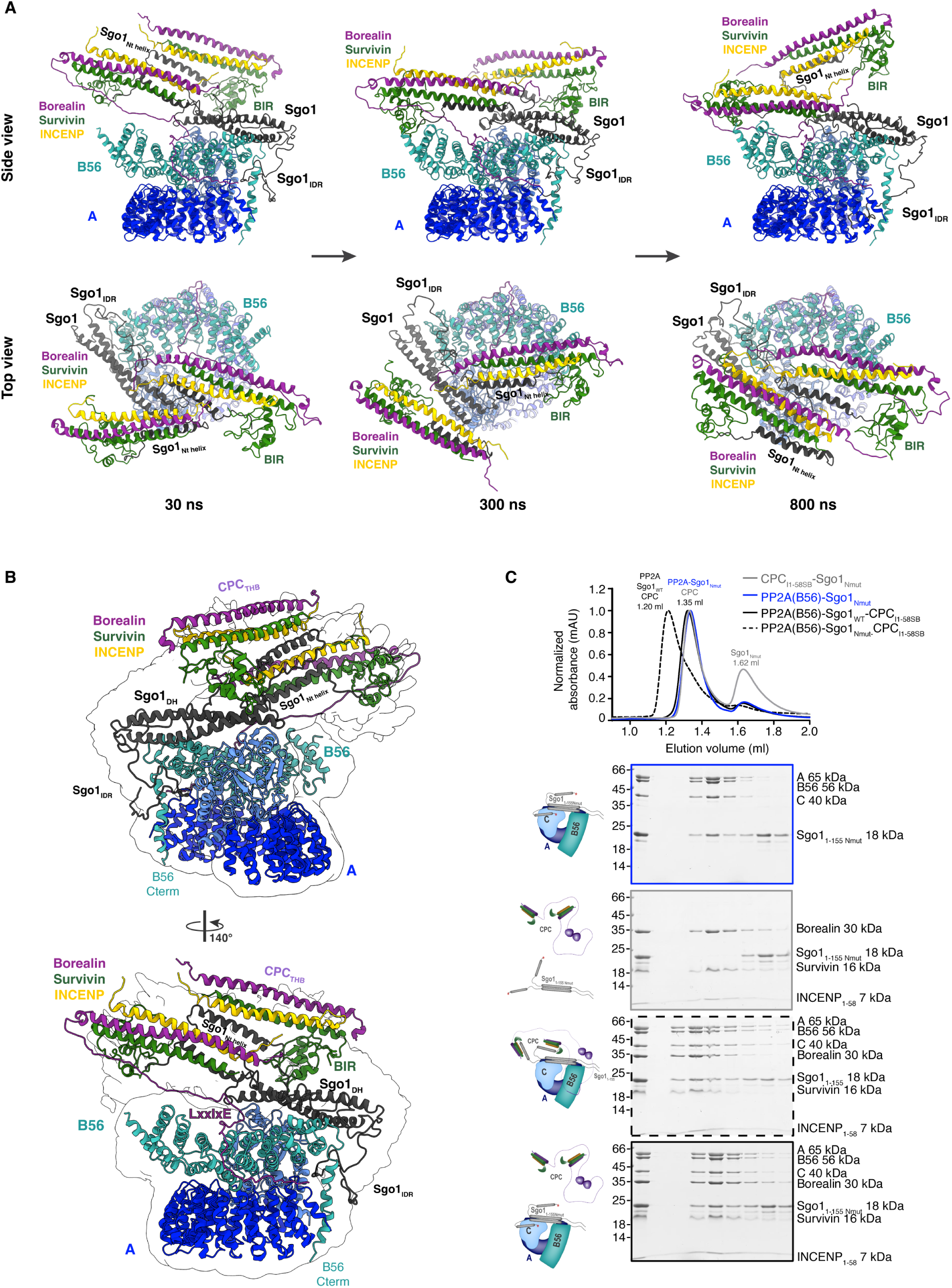
Overall architecture of PP2A(B56)-Sgo1-CPC_I1-58SB_ using computational modeling and molecular dynamics simulation. **(A)** Snapshots of the MD trajectories of the PP2A(B56)-Sgo1-CPC_I1-58SB_ complex (two views) at 30 ns, 300 ns and 800 ns of the simulation. **(B)** Selected frame of the MD simulation for the PP2A(B56)-Sgo1-CPC_I1-58SB_ complex fitted into the gaussian blurred ‘consensus’ map, in two orientations. **(C)** SEC profiles and corresponding SDS-PAGEs of PP2A(B56) (blue), PP2A(B56)-Sgo1_Nmut_ (grey), PP2A(B56)-Sgo1-CPC_I1-58SB_ (dashed black) and PP2A(B56)-Sgo1_Nmut_-CPC_I1-58SB_ (solid black).

**Figure S8:**
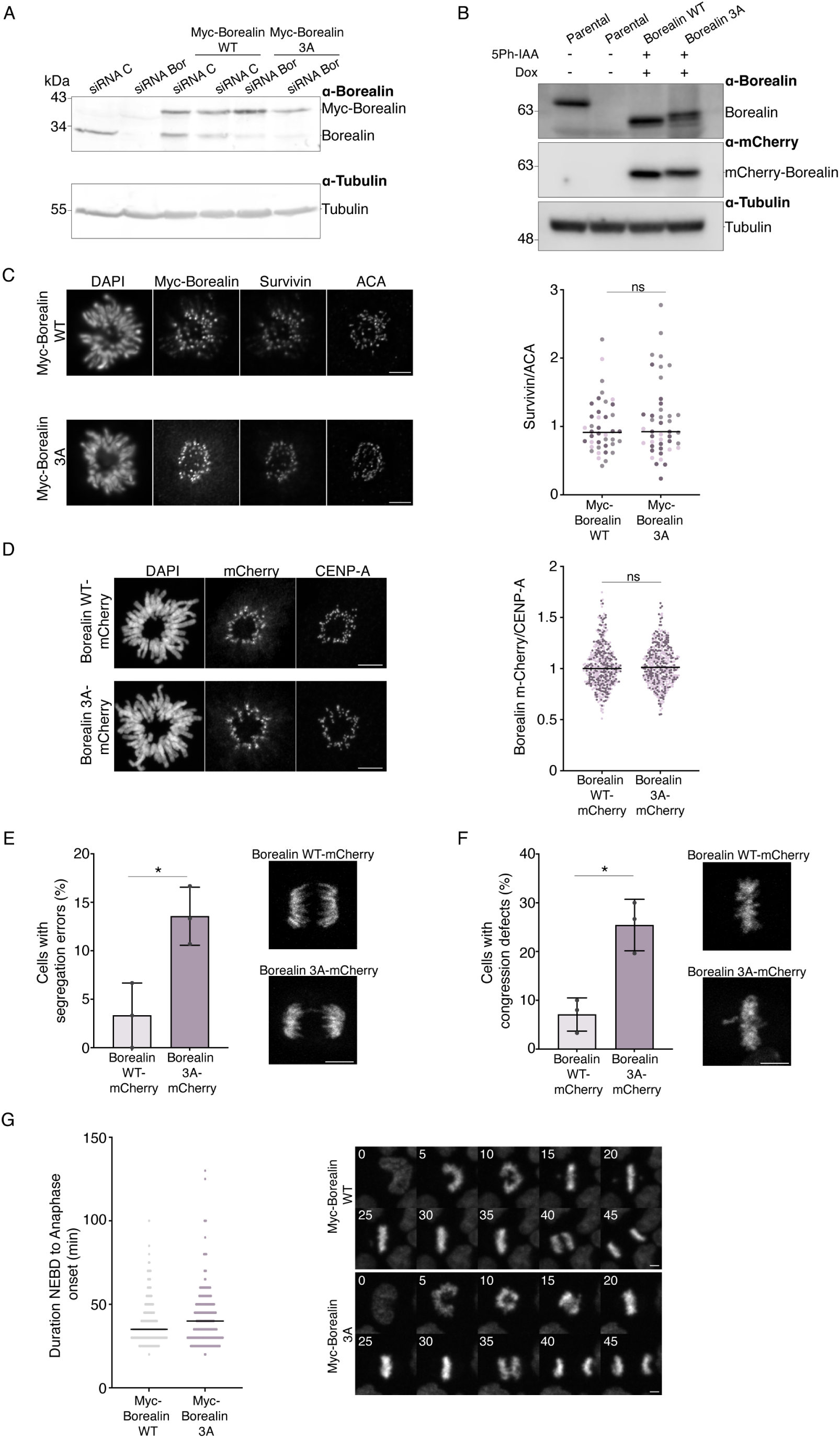
Loss of PP2A(B56)-CPC interaction leads to chromosome segregation defects. **(A, B)** Western blot analysis of Borealin depletion and expression of WT or 3A Borealin constructs in HeLa Kyoto (A) and hTERT-RPE1 (B) cells. **(C)** Quantification of centromeric Survivin levels, normalized to ACA intensity, in HeLa Kyoto cells following Borealin depletion and rescue with Myc-Borealin_WT_ or Myc-Borealin_3A_. DNA was stained with DAPI. Scale bar, 5 μm. Lines indicate medians. Data represent three independent experiments (≥40 cells); Mann– Whitney U test; ns, not significant. **(D)** Representative images of Borealin-mCherry and CENP-A localization in prometaphase RPE1 cells expressing Borealin-mCherry WT or 3A following endogenous Borealin replacement. DNA was stained with DAPI. Scale bar, 10 μm. Right, quantification of Borealin-mCherry intensity normalized to CENP-A. A total of 210 kinetochores from seven cells per condition were analysed. Each dot represents one kinetochore. Statistical analysis used mean values from 30 kinetochores per cell (unpaired two-tailed t-test). The experiment was repeated twice; ns, not significant. **(E, F)** Incidence of chromosome missegregation (E) and congression defects (F) in RPE1 cells expressing Borealin-mCherry WT or 3A. Data represent three independent experiments (≥30 cells each); bars indicate mean ± SD. Paired two-tailed t-test; *, P ≤ 0.05. Scale bar, 10 μm. **(G)** Representative live-cell images (5-min intervals) and quantification of mitotic progression in HeLa cells expressing Myc-Borealin_WT_ (pink) or Myc-Borealin_3A_ (purple). NEBD-to-anaphase timing was measured in ≥205 cells across three biological replicates. Each dot represents one cell; lines indicate medians. DNA was visualized using SPY650-DNA. Scale bar, 5 μm.

**Supplementary Table S1:** Cryo-EM data collection, refinement and validation statistics.

|  | **PP2A(B56)-Sgo1** high-resolution map | **PP2A(B56)-Sgo1** consensus map | **PP2A(B56)-CPC<sub>11-58SB</sub>** high-resolution map | **PP2A(B56)-CPC<sub>11-58SB</sub>** Borealin loop and DD map | **PP2A(B56)-Sgo1-CPC<sub>11-58SB</sub>** high-resolution map | **PP2A(B56)-Sgo1-CPC<sub>11-58SB</sub>** consensus map | **PP2A(B56)-Sgo1-CPC<sub>11-58SB</sub>** Borealin loop map |
|---|---|---|---|---|---|---|---|
| **PDB** | 50OA | - | 50NZ | - | 50OC | - | - |
| **EMDB** | EMD-100801 | EMD-100801 | EMD-100800 | EMD-100800 | EMD-100802 | EMD-100802 | EMD-100802 |
| **Data collection and processing** |  |  |  |  |  |  |  |
| *Microscope/detector* | Titan Krios / Falcon4i | Titan Krios / Falcon4i | Titan Krios / Falcon4i | Titan Krios / Falcon4i | Titan Krios / Falcon4i | Titan Krios / Falcon4i | Titan Krios / Falcon4i |
| *Voltage (ke<sup>-</sup>V)* | 300 | 300 | 300 | 300 | 300 | 300 | 300 |
| *Magnification (x)* | 165k | 165k | 165k | 165k | 165k | 165k | 165k |
| *Defocus* | -0.5 µm to -2.6 µm (step size 0.3 µm) | -0.5 µm to -2.6 µm (step size 0.3 µm) | -0.5 µm to -2.6 µm (step size 0.3 µm) | -0.5 µm to -2.6 µm (step size 0.3 µm) | -0.5 µm to -2.6 µm (step size 0.3 µm) | -0.5 µm to -2.6 µm (step size 0.3 µm) | -0.5 µm to -2.6 µm (step size 0.3 µm) |
| *Pixel size (Å)* | 0.727 | 0.727 | 0.727 | 0.727 | 0.727 | 0.727 | 0.727 |
| *Total dose (e<sup>-</sup>/Å<sup>2</sup>)* | 40 | 40 | 40 | 40 | 40 | 40 | 40 |
| *Number of micrographs* | 10,182 | 10,182 | 10,356 | 10,356 | 16,787 | 16,787 | 16,787 |
| *Symmetry imposed* | C1 | C1 | C1 | C1 | C1 | C1 | C1 |
| *Initial number of particles* | 273,925 | 273,925 | 153,131 | 153,131 | 388,545 | 388,545 | 388,545 |
| *Final number of particles* | 79,131 | 27,278 | 71,070 | 16,872 | 121,329 | 121,784 | 31,522 |
| *Map resolution (Å) (FSC 0.143)* | 3.11 | 3.78 | 3.05 | 3.51 | 2.82 | 2.89 | 3.14 |
| **Refinement** |  |  |  |  |  |  |  |
| *Initial model used* | AF3 | - | AF3 | - | AF3 | - | - |
| *Map CC* | 0.84 | - | 0.83 | - | 0.86 | - | - |
| *Model composition* |  |  |  |  |  |  |  |
| *Non-hydrogen atoms* | 11219 | - | 10382 | - | 11286 | - | - |
| *Protein residues* | 1403 | - | 1300 | - | 1411 | - | - |
| *Ligands* | MN:2 | - | MN:2 | - | MN:2 | - | - |
| *Mean B factor (Å<sup>2</sup>)* |  |  |  |  |  |  |  |
| *Protein* | 81.19 | - | 98.64 | - | 60.30 | - | - |
| *Ligand* | 73.31 | - | 37.63 | - | 42.97 | - | - |
| *RMS deviation* |  |  |  |  |  |  |  |
| *Bond length (Å)* | 0.004 | - | 0.004 | - | 0.005 | - | - |
| *Bond angle ( ° )* | 1.020 | - | 1.014 | - | 1.002 | - | - |
| *Validation* |  |  |  |  |  |  |  |
| *All atom clash score* | 7.04 | - | 5.70 | - | 5.81 | - | - |
| *Rotamer outliers (%)* | 2.42 | - | 1.75 | - | 2.25 | - | - |
| *MolProbity score* | 1.70 | - | 1.50 | - | 1.58 | - | - |
| *Ramachandran plot* |  |  |  |  |  |  |  |
| *Favored (%)* | 97.92 | - | 97.99 | - | 98.28 | - | - |
| *Allowed (%)* | 2.08 | - | 2.01 | - | 1.72 | - | - |
| *Outliers (%)* | 0 | - | 0 | - | 0 | - | - |

## Notes

### Competing Interest Statement

The authors have declared no competing interest.

